# A multi-region recurrent circuit for evidence accumulation in rats

**DOI:** 10.1101/2024.07.08.602544

**Authors:** Diksha Gupta, Charles D. Kopec, Adrian G. Bondy, Thomas Z. Luo, Verity A. Elliott, Carlos D. Brody

## Abstract

Decision-making based on noisy evidence requires accumulating evidence and categorizing it to form a choice. Here we evaluate a proposed feedforward and modular mapping of this process in rats: evidence accumulated in anterodorsal striatum (ADS) is categorized in prefrontal cortex (frontal orienting fields, FOF). Contrary to this, we show that both regions appear to be indistinguishable in their encoding/decoding of accumulator value and communicate this information bidirectionally. Consistent with a role for FOF in accumulation, silencing FOF to ADS projections impacted behavior throughout the accumulation period, even while nonselective FOF silencing did not. We synthesize these findings into a multi-region recurrent neural network trained with a novel approach. In-silico experiments reveal that multiple scales of recurrence in the cortico-striatal circuit rescue computation upon nonselective FOF perturbations. These results suggest that ADS and FOF accumulate evidence in a recurrent and distributed manner, yielding redundant representations and robustness to certain perturbations.

## Introduction

Decision-making based on noisy perceptual evidence is a core cognitive process that is often conceptualized as a sequence of two sub-computations: gradual accumulation of evidence, followed by thresholding to commit to a categorical choice. Both theoretical^1,2,3,4^ and empirical^5,6,7,8^ studies of the neural implementation of this process have proposed candidate circuits with feedforward information flow, by mapping the two sub-computations onto distinct brain areas. Such a modular implementation offers several adaptive advantages^9^; for instance it allows for easy dynamic modulation of decision thresholds based on behavioral demands while leaving the accumulation process unchanged^10,11,12,3,13^.

In rats, the anterior dorsal striatum (ADS) and the frontal orienting fields (FOF) have been mapped onto these two theoretically defined sub-computations. ADS is known to represent the graded accumulated evidence and is causally required throughout the accumulation process^14^, whereas FOF is known to represent the categorical choice and is causally necessary only at the very end of accumulation period when the graded evidence needs to be thresholded^15,16,17,18^. These observations provide an intriguing, yet untested, neural implementation of the decision process, in which ADS and FOF form a feedforward functional hierarchy - with evidence accumulated in ADS being thresholded in FOF.

While appealing, this proposal is surprising in several ways. First, the fact that FOF projects monosynaptically to ADS, but ADS does not project directly back to FOF, implies that this proposed implementation must map onto a multisynaptic pathway in which signals from ADS must be relayed through other basal ganglia nuclei, thalamus and/or superior colliculus before they reach FOF. Furthermore, FOF is one of the main cortical inputs to ADS, but this salient FOF → ADS projection plays no role in the proposed implementation. Anatomically connected brain areas can selectively communicate information to their downstream targets^19,20^, so it is certainly possible that the information flow during evidence accumulation renders the monosynaptic projection from FOF to ADS non-functional and only involves the multisynaptic pathway from ADS to FOF. However, the substantial heterogeneity in neural encoding within FOF, with neurons representing both graded evidence and categorical choice^16^, along with FOF’s anatomical placement, strongly suggests that FOF might have a role in gradual accumulation as well.

Second, some recent studies challenge the interpretation of transient unilateral optogenetic perturbations of FOF^16^, which had previously provided compelling evidence for confining FOF’s role to the thresholding epoch of decision-making. It has been observed that, unlike its response to bilateral perturbations, the cortex can be robust to transient unilateral perturbations – with the network recovering through information from the other hemisphere^21^. Since the network has enough time to recover early in the accumulation period but not towards the end, this interhemispheric compensation might underlie the lack of impairment observed during FOF’s inactivations early in the decision process, obscuring its role. Furthermore, whole-region non-specific perturbations have been shown to conceal effects that can be revealed by more specific perturbations that target a particular projection^22,23,24^. This suggests that the previous unilateral inactivation studies may have missed aspects of FOF’s involvement in the decision process by not using projection-specific and/or bilateral manipulations.

Therefore, it is crucial to directly evaluate the hypothesized feedforward and modular mapping of the decision process onto ADS and FOF. Testing this would require recording activity from both areas simultaneously^25,26^ to determine if the distinct representations as well as the transformations and latencies expected from feedforward communication, are present across the two areas. Such a comparison would also control for slight differences in cognitive strategies or training histories between subjects that can vary neural representations and dynamics^27,28,29,30,31^. Additionally, confirming that the feedback projection under this hypothesis i.e. the FOF → ADS projection plays no causal role in evidence accumulation would add substantial support in favor of this hypothesis. Similarly, demonstrating undisrupted evidence accumulation during bilateral manipulations of FOF activity would address the aforementioned confounds and further validate that FOF’s role is confined to the thresholding sub-computation.

Here, we follow this logic and apply simultaneous multi-region population recordings, bilateral, and projection-specific perturbations in rats performing an evidence accumulation task. The resulting data provide evidence against the modular, feedforward hypothesis in which evidence accumulated in ADS is thresholded in FOF. We find that ADS does not have privileged information about evidence, rather both regions carry similar amounts of information that evolves at comparable timescales. Further, simultaneous population recordings let us define the feedforward (ADS → FOF) and feedback (FOF → ADS) communication subspaces under this hypothesis. We find that evidence can be substantially decoded from both these subspaces, in contradiction to the hypothesized information flow. Moreover, with projection-specific inactivations we show that this “feedback” communication is necessary during the accumulation process for unimpaired decision-making.

Therefore, we revise the feedforward hypothesis and advocate for an alternate distributed implementation of the evidence accumulation process in the cortico-striatal circuitry. We instantiate this proposal using a multi-region recurrent neural network model with biological constraints on connectivity. The model successfully reproduces empirical representational redundancies and effects of held-out perturbations, when trained with a novel approach of capturing patterns of animal behavior on optogenetic perturbation trials alongside control trials. Finally, leveraging the RNN’s observability and controllability, we predict that recovery dynamics are at play during non-selective silencing of FOF neurons and perform a number of in-silico experiments to understand the contributions of different projections to this phenomenon. We show that recovery is supported by multiple scales of recurrence - including self-recurrence, inter-hemispheric recurrence and striato-cortical recurrence. Taken together, our approach of using projection-specific perturbations in conjunction with multi-region RNN modeling offers a useful method to decipher the contributions of distributed and recurrently connected networks of brain regions to cognitive processes.

## Results

In order to test whether the hypothesized transformation between ADS and FOF holds on single trials (Figure 1B-C), we recorded simultaneous population activity from ADS and FOF of rats trained to perform a previously developed decision-making task^32^ (Fig 1A). This task requires accumulation of auditory evidence over hundreds of milliseconds for good performance. On any given trial, rats are presented with two streams of randomly timed auditory clicks, one played from a speaker to their left and the other from a speaker to their right. At the end of the stimulus, rats are rewarded with a drop of water for correctly reporting the side which played the greater number of clicks.

**Figure 1.**
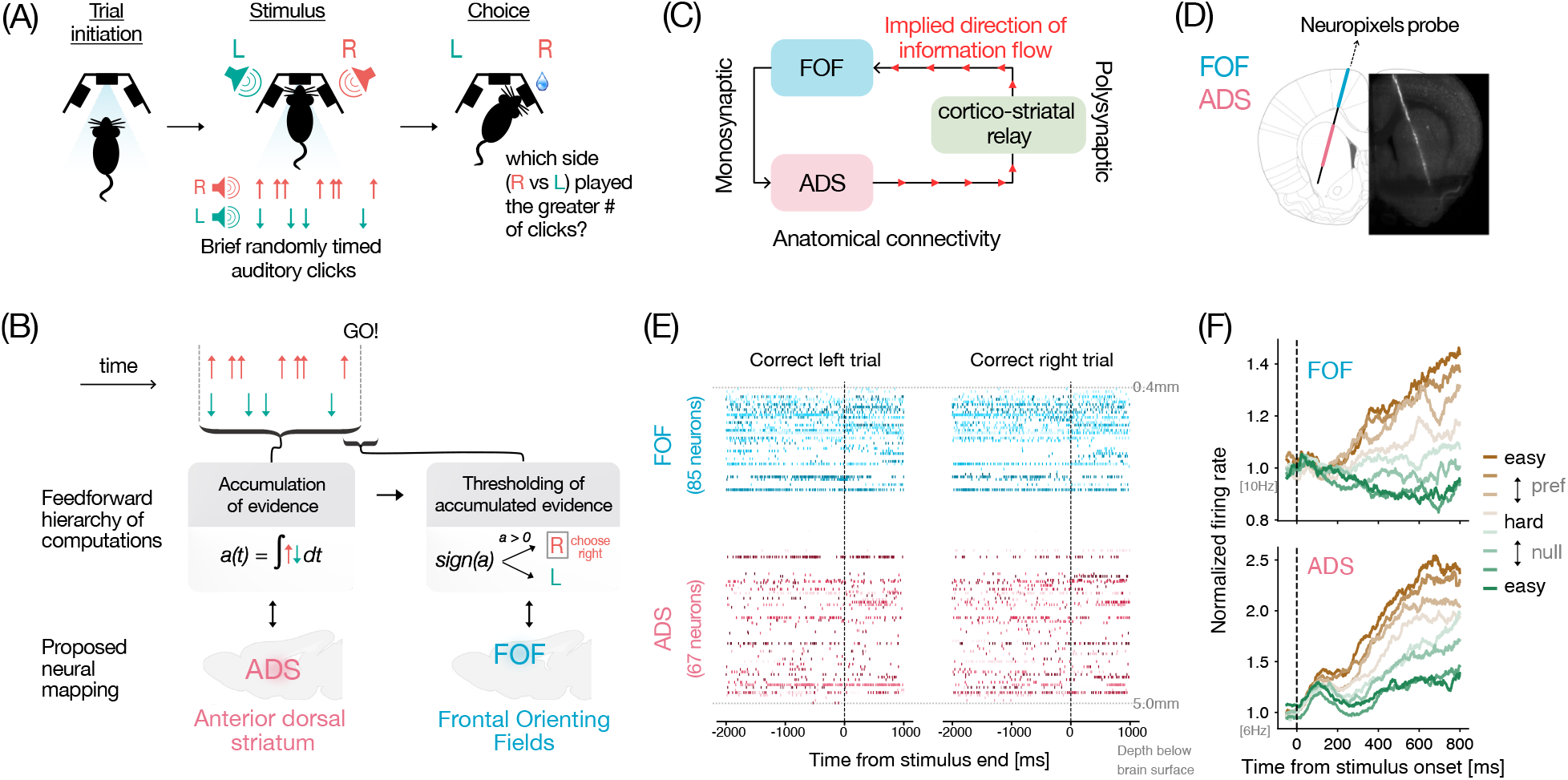
Experimental setup and simultaneous recordings from FOF and ADS. **(A)** Schematic of the events in the rat evidence accumulation task (adapted from Brunton et al. 2013). From left to right: rats initiate a trial by poking their nose into the center port, two streams of randomly timed auditory clicks play from speakers on their left and right side, rats are rewarded for choosing the side with the greater number of clicks. **(B)** Schematic showing proposed implementation of the decision-making process^16,14,38^. The anterior dorsal part of striatum (ADS, pink) is thought to be involved in gradual accumulation of sensory evidence and represents this accumulated value “*a*” in a graded fashion^14^. The accumulated evidence is thought to be thresholded, and represented in categorical form in the frontal orienting fields (FOF, blue), a part of the rat secondary motor cortex^16,15,17^. **(C)** The pathway from ADS to FOF involves a multi-synaptic pathway (grey arrows through olive-green regions) through basal ganglia (BG) nuclei, the thalamus and the superior colliculus (SC) that could potentially support this transformation. **(D)** Schematic showing neuropixels targeting (AP 1.9, ML 1.15 at a 15 degrees angle in the coronal plane) to record simultaneously from FOF and ADS, overlaid on histology from an example rat. Brain atlas was adapted from Paxinos and Watson, 2006 **(E)** Spike rasters showing simultaneous neural activity recorded with a neuropixels probe from two example correct trials (left and right) from one session. Neurons are arranged in order of their estimated depth from the brain surface. Aqua blue shades represent FOF neurons and pink shades represent ADS neurons. **(F)** Average FOF (top) and ADS (bottom) population responses during stimulus period for different strengths of sensory evidence. Brown and green colors correspond to stimuli from preferred and non-preferred sides of the neurons, respectively. Darker hues correspond to easier trials and greyish hues correspond to harder trials. Both areas show stimulus strength dependent ramping activity. (FOF n = 219 neurons, ADS n = 235 active side-selective neurons from 5 rats).

While rats (n = 5) performed this task, we used Neuropixels probes^33,34^ to record neural activity from FOF and ADS simultaneously (Fig 1D-E). We recorded from 765 FOF and 1583 ADS neurons across 12 sessions. Cells with a mean firing rate of at least 1Hz during the stimulus period were considered active and included for further analysis (462 FOF, 559 ADS neurons). In both FOF and ADS, we found that neurons that significantly modulated their activity in a side-selective manner on average exhibited firing rates that ramped upwards for stimuli favoring their preferred side with a slope proportional to the stimulus strength (Fig 1F). These trial-averaged stimulus-dependent ramping responses have long been interpreted as a neural correlate of the evidence accumulation process^5^ and are consistent with previous reports in rats from FOF^16^ and ADS^14^, and analogous regions in primates (FEF^35^ and caudate^36^). However, both graded and categorical representations of accumulated evidence can give rise to such ramping responses^37,16^, hence to distinguish between these possibilities we characterized the encoding and decoding profiles of task-relevant variables in the two regions with more directed analysis.

### FOF and ADS carry redundant bidirectionally communicated task-related information

The feedforward functional hypothesis posits that ADS has graded encoding of evidence that is relayed to FOF where this evidence is thresholded and the categorical choice is represented. Therefore, the hypothesis predicts that evidence can be decoded from ADS earlier and with higher accuracy compared to FOF.

To test the validity of this hypothesis, first we sought to compare the differences in the extent of graded encoding between FOF and ADS. For this we use the method developed in Hanks 2015^16^. This method computes the accumulator ‘tuning curves’ i.e. maps of how firing rates of neurons vary as a function of the accumulator value. To do so, it leverages the behavioral model from Brunton 2013^32^ to obtain trial-by-trial, moment-by-moment estimates of the subject’s accumulator value and relates it to the simultaneously observed firing rates of neurons. Previous application of this method had found that tuning curves in FOF had high slopes, i.e. were more step-like or categorical, with accumulator values on one side of the decision boundary giving rise to one cluster of firing rates, and the values on the other side of the decision boundary giving rise to another cluster^16^. On the other hand, previous application of this method in ADS had found a prevalence of neurons whose firing rates had lower slopes i.e. varied smoothly with the graded value of the accumulator^14^. Here, simultaneous recordings allowed us to perform comparisons controlled for inter-subject variability.

When applying this analysis to side-selective neurons in our data, we found examples of both graded and categorically tuned neurons in FOF as well as ADS (Fig 2A). Surprisingly, we found no significant differences between FOF and ADS (Fig 2B, P = 0.13 Mann-Whitney U test). If anything, the median slope was slightly higher in ADS (0.32 ± 0.25, median ± std) compared to FOF (0.29 ± 0.22). Contrary to expectations from previous studies, these results demonstrate that when neural measurements from the two areas are made in the same subjects, the two regions do not appear to differ in their encoding of the accumulator variable.

**Figure 2.**
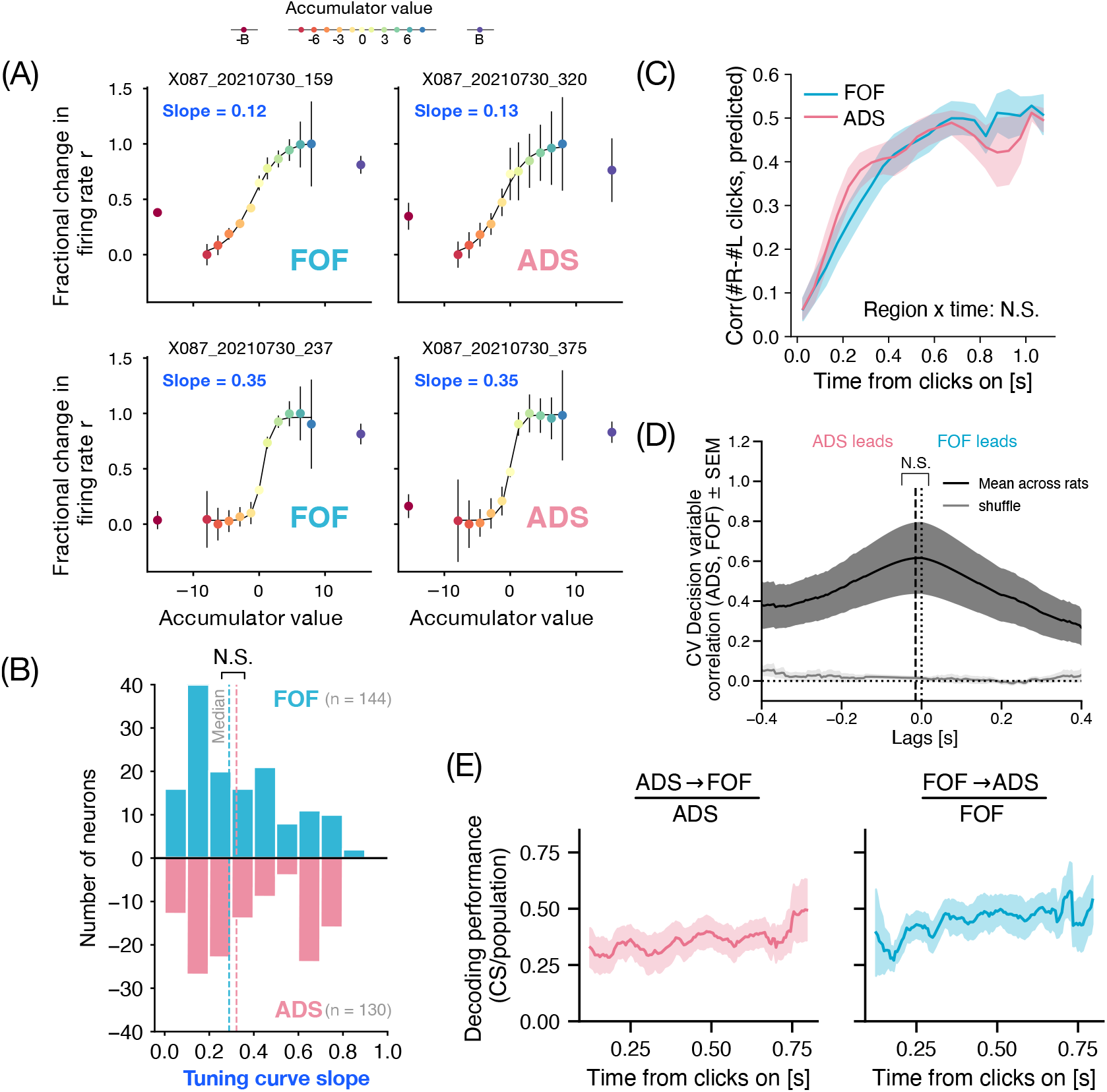
FOF and ADS carry redundant bidirectionally communicated task-related information. **(A)** Tuning to the behaviorally inferred accumulator variable of two example neurons from FOF (left) and ADS (right) each. (Top panel) neuron exhibits a graded response to the accumulator variable, signified by the shallow slope of its tuning curve. (Bottom panel) exhibits a sharp dependence on the sign of the accumulator variable, reflected in the steeper slope of its tuning curve. **(B)** Histogram of tuning curve slopes in FOF (aqua blue; n = 144) and ADS (pink; n = 130). Dashed lines indicate the median slopes for the two populations (FOF = 0.29 ± 0.22; ADS = 0.32 ± 0.25, median ± std). The two populations have similar distribution of tuning to the accumulator variable, with neurons representing the accumulator value in both graded (smaller slopes) and categorical (larger slopes) fashion. The medians of the two populations were not significantly different (P = 0.13, Mann Whitney U test). **(C)** Stimulus decoding performance (mean ± sem) of the linear decoder on held-out time points as a function of time from stimulus onset (n = 12 sessions). Linear decoders were fit independently to the two populations (FOF: aqua blue; ADS: pink) while controlling for the number of neurons from the two regions. Performance is measured as the correlation between the actual trajectory of difference in number of clicks and the prediction from linear decoding of neural responses. Decoding performance and its timecourse did not significantly differ between FOF and ADS (P = 0.36, two-way RM ANOVA) **(D)** Time-lagged cross-correlogram between decision-variable trajectories decoded from FOF and ADS, showing the correlation coefficient (y-axis) at different time lags (x-axis: positive (negative) values represent FOF trajectories leading (lagging) ADS). Solid black line indicates mean across sessions and rats, with dark-gray shaded region representing S.E.M. Light gray region indicates a shuffled control, computed by cross-correlating FOF activity with ADS activity on trials with shuffled (rather than matched) identities. The cross-correlogram is symmetric with a wide peak at a latency not significantly different from zero (mean latency ± sem = −0.01 ± 0.02s; P = 0.47 one-sided t-test, n = 12), hence showing no strong evidence of any lead-lag relationships. **(E)** Time-course of stimulus decoding accuracy present in the ADS → FOF (pink) and FOF → ADS (blue) communication subspaces (CSs), plotted as a fraction of stimulus decoding accuracy from the entire region’s population activity. A fraction of 1 indicates that the decoding from CS is just as good as the entire population. Stimulus information is present in both regions’ CS and increases over the course of the trial, with FOF sharing a slightly higher fraction of its information in the CS.

Next, we compared the decodability of information about the stimulus and other task variables in the population activity of FOF and ADS. Despite the similarity in trial-averaged tuning to the accumulator variable (Fig 2B), it is still possible that this information is decodable earlier in ADS, and/or with higher accuracy. For each region, we trained separate linear decoders to predict the cumulative difference in the number of right and left clicks at each time point during the trial (Supp Fig 1A). Interestingly, both regions showed comparable decoding performance over time, as measured by the correlation between the true and predicted cumulative stimulus on held out time points, with no significant differences between the regions (Fig 2C; two way repeated measures ANOVA, P = 0.93 for region and P = 0.36 for region x time). These results did not change when choice information was controlled for, by separately decoding the stimulus on left and right choice trials (Supp Fig 1B). The results were similar also when instead of the veridical stimulus, the mean accumulated evidence inferred with the behavioral model was decoded from the two populations (not shown).

Additionally, we trained logistic decoders to predict other binary task-relevant variables such as choice and trial-history from the two regions. The two regions showed high decoding of choice (Supp Fig 1C) but once again with no significant differences in decoding performance over time (two way repeated measures ANOVA, P = 0.90 for region and P = 0.54 for region x time). The past trial’s choice could be decoded well into the current trial with substantial accuracy from both regions (Supp Fig 1D), however there were no significant differences between the two regions (two way repeated measures ANOVA, P = 0.13 for region and P = 0.37 for region x time). Altogether, these results show that, at least under the analyses carried out here, the two regions have indistinguishable representations of task relevant variables, leading us to believe that their involvement in decision-making may be strongly related.

Our conclusions about the simultaneous evolution of evidence-related representations in FOF and ADS are also supported by an independent analysis that decodes the decision variable from neural population activity. The distance of a population firing rate vector from the decision hyperplane of a logistic decoder that is trained to predict choice, reflects the model’s prediction confidence. This neurally inferred variable, known as the decision variable (DV), estimates the animal’s internal decision variable (log odds of making a choice) capturing the influence of sensory evidence and other choice determinants^39,21,40^. This complements our previous analyses by relaxing assumptions of the behavioral model and not assuming a fixed stimulus encoding lag in the two regions. We sought to examine if these neurally inferred DVs from the two regions exhibit any systematic lead-lag relationships in their evolution, in order to infer the dominant direction of task-related information flow within the frontal cortico-striatal circuitry. We used the logistic decoder trained to predict the eventual choice (from Supp Fig 1C) and estimated the DV on single trials from the two regions (Supp Fig 2A). DVs from FOF and ADS showed good correspondence with each other and with the eventual true choice of the animal on both correct and error trials (Supp Fig 2B). Averaged across trials of different difficulties, DV values (unsigned) increased over the trial, with the rate of increase dependent on the strength of sensory evidence – exhibiting the classic ramping signature expected from evidence accumulation (Supp Fig 2C). Consistent with our previous analyses, we did not find any lead-lag relationship in the evolution of DV in the two regions (Fig 2D). While the DVs showed high correlations at short latencies on average, the cross-correlogram of these DVs averaged over rats and sessions had a peak at a latency not significantly different from zero (mean latency ± sem = −0.01 ± 0.02s; P = 0.47 one-sided t-test, n = 12). This is consistent with no dominant feedforward or feedback relationship^41^. We conducted extensive simulations to validate our ability to infer lead-lag relationships in evolution of DV across varying number of simultaneously recorded neurons, trials and interaction lags (Supp Fig 3).

**Figure 3.**
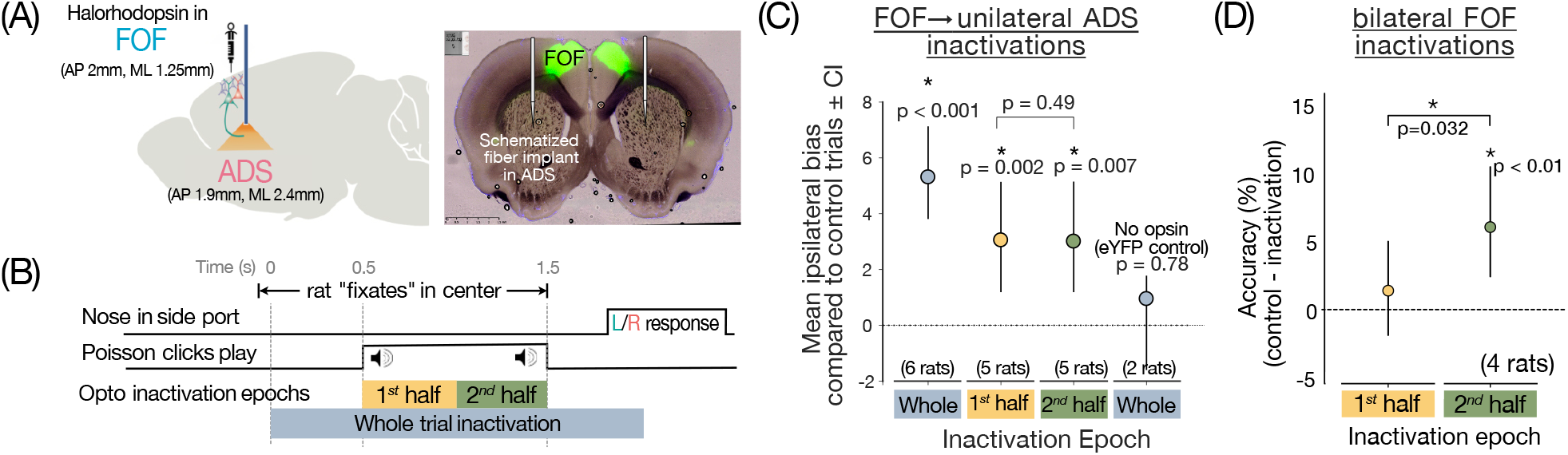
Silencing FOF inputs to ADS disrupts decision-making. **(A)** Silencing FOF axon terminals in ADS: experimental setup (Left) Schematic showing viral delivery of Halorhodophsin (eNpHR3.0) to FOF and sharp optical fiber implant in ADS. Virus was injected bilaterally and 594mm wavelength laser (25-33mW) was delivered unilaterally through the fiber optic cable on 25% of trials to silence FOF axon terminals in ADS. (Right) Example histology from a rat expressing eNpHR3.0 bilaterally in FOF. **(B)** Time-line of task events showing perturbation epochs: FOF axon terminals in ADS were perturbed during one of three epochs: *Whole-trial* (grey bar) started as soon as the rat entered the center port and terminated 500ms after the stimulus ended. *Early* (yellow) spanned the first half of a 1s long stimulus and *late* (green) spanned the second half of a 1s long stimulus. It is predicted that if the FOF→ADS projection is involved in conversion of the graded evidence to a categorical choice, then the inactivation should have an effect on behavior only at the end of the stimulus presentation i.e. during the late epoch. Whereas, if the projection is involved in evidence accumulation, a process that occurs throughout the stimulus presentation, inactivations during both early and late epochs should have an effect. **(C)** The ipsilateral bias, or the difference in percentage of trials the rat went to the side ipsilateral to the laser on inactivation v.s. control trials plotted for different inactivation epochs. Circles represent the mean ipsilateral bias and error bars represent 95% bootstrap confidence intervals. Whole-trial inactivations (6rats, 10 hemispheres) in rats injected with NpHR produced a significant impairment in behavior (P<0.001, non-parametric bootstrap test) and no significant effects in the eYFP controls (P=0.78) (2rats, 3 hemispheres). Early and late inactivations (5 rats, 8 hemi-spheres) also produced significant impairments (P<0.005 non-parametric bootstrap test) with no significant differences between the effects of the two epochs (P=0.49) suggesting that FOF-ADS projections are involved throughout the accumulation process. **(D)** Difference in accuracy between control and inactivation trials, plotted for temporally specific bilateral inactivations of FOF in early (yellow) or late (green) stimulus epochs. Circles represent mean accuracy difference across 4 rats, error bars represent 95% bootstrap confidence intervals. Only inactivations in the late epoch produce significant reductions in accuracy (p<0.01, nonparametric bootstrap test), differing significantly from inactivations in early epochs (p<0.032, nonparametric bootstrap test) in which there is no significant impairment.

Next, we probed the content of the interaction between FOF and ADS during decision-making more generally, rather than restricting ourselves to activity along the decision hyperplane. To measure such interactions we used a generalized linear model with coupling filters across the two populations (similar to Pillow 2008^42^) in addition to temporally-extended kernels for left and right clicks and other major task events. To overcome the explosion in the number of parameters with increasing numbers of cells, instead of modeling pairwise coupling terms, we projected the activity of the regressor population onto a (learnt) lower dimensional subspace, over which the coupling filters were learnt (Methods; Supp Fig 4A). We estimated the number of such latent factors required by evaluating the cross-validated log-likelihood of models fit with varying dimensions (Supp Fig 4B). The resultant model captured a high percentage of variance in average firing rates for most of the neurons in both areas (Supp Fig 4C). We compared the information content of activity in FOF → ADS and ADS → FOF communication subspaces (CS) to the whole population activity space by measuring the decoding accuracy of stimulus (Fig 2E). Remarkably, we found that a sizeable fraction of stimulus information in the two regions could also be decoded from the CS and this fraction increased over the course of the trial, with FOF sharing a slightly higher fraction of its information in its CS 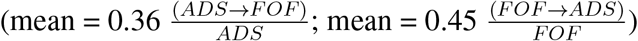. Altogether, these results indicate that the representations of evidence in the two regions exist in the space that captures the shared trial-by-trial noise fluctuations and therefore are likely communicated between the two.

**Figure 4.**
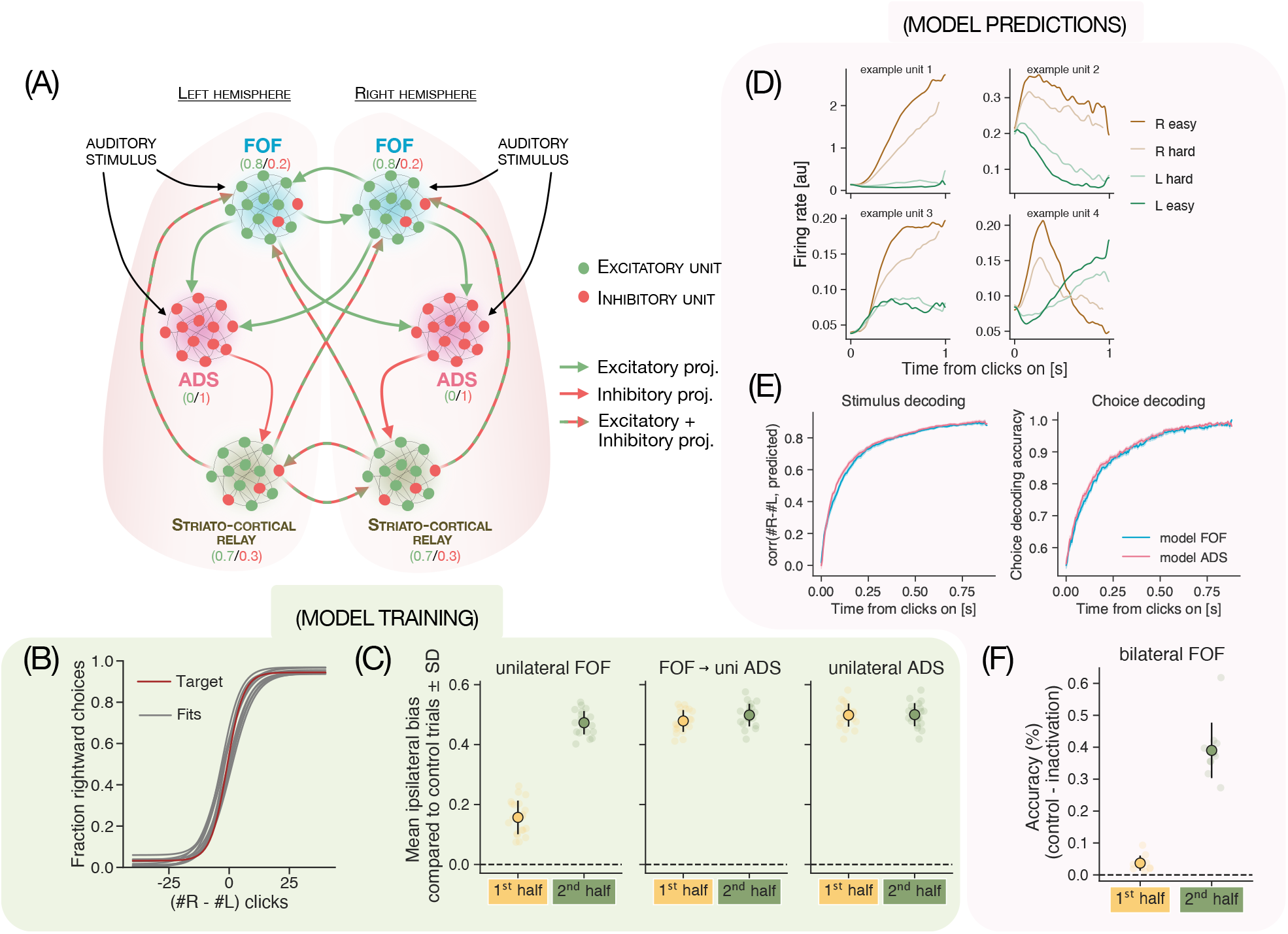
Multiregion recurrent neural network model successfully recapitulates empirical representational and robustness properties. **(A)** Schematized model architecture. The model has two modules that represent the two brain hemispheres. Within each module are three submodules that correspond to FOF, ADS and the striato-cortical relay. Inhibitory projections and units are depicted in red, and excitatory projections and units are depicted in green. All units in the model followed Dale’s law. The inputs were sent to all excitatory units in FOF and ADS and the outputs were read out from all units in the two hemispheres. **(B)** Psychometric curve for RNN model-produced choices (grey; n = 10) overlayed on target psychometric produced by the target choices from the Brunton model (red). **(C)** Average bias towards the ipsilateral choices exhibited upon unilateral (uni) inactivations of FOF, FOF→ADS projections and ADS (left to right) during either the first or second half of the stimulus. Bars represent the mean ± CI across 10 trained networks, each dot represents results from inactivations in one of the hemispheres of the trained networks. The network recapitulates empirical data and shows high ipsilateral bias upon inactivation of FOF in the 2nd half compared to the 1st half. FOF→ADS and ADS inactivations produce ipsilateral bias both in the 1st and 2nd halves, resembling rat data. **(D)** PSTHs (aligned to stimulus onset, conditioned on stimulus difficulty) for example units from the model. Model units show variable timecourses and diversity in when the strength of evidence is encoded. **(E)** Model FOF and ADS have similar timecourse of stimulus and choice decoding (Left) Correlation between the true cumulative difference in number of right and left clicks at any time point during the trial and that predicted by a linear decoder trained on either model FOF activity (aqua blue) or model ADS activity (pink), as a function of time from stimulus onset. (Right) Accuracy of a logistic decoder trained to predict the eventual choice from responses of either model FOF (aqua blue) or ADS (pink) during the stimulus. **(F)** Model successfully predicts response to novel perturbations: difference in accuracy between model’s choices on control trials and trials in which model FOF was inactivated bilaterally either during the first (orange) or the second (green) half of the stimulus. Bars represent the mean ± CI across 10 trained networks, dots represent the difference for individual networks.

### Silencing FOF inputs to ADS disrupts decision-making

From the analysis of simultaneous activity of FOF and ADS we observed that the two regions have remarkably similar encoding and decoding of decision-relevant variables, this information is shared in their communication subspaces, and that there is no discernible lead-lag relationship between the representations of the evolving decision variable in the two regions. This redundancy could indicate that these signals are inherited by both FOF and ADS from a third (unknown) brain region. However, given the causal involvement of these regions during decision-making, a more interesting and likely possibility is that this redundancy reflects ongoing recurrent interactions^41^ between FOF and ADS. Are these interactions necessary for making decisions? Establishing whether or not these interactions are important would put strong constraints on the network mechanisms that underlie decision-making and help further assess the feedforward, modular hypothesis.

We sought to probe this question in the experimentally tractable monosynaptic projection from FOF to ADS using unilateral optogenetic inactivations. We injected the inhibitory opsin, halorhodopsin (AAV5-CaMKII*α*-eNpHR3.0-eYFP) bilaterally in FOF and delivered light (25-33mW, 594nm) via a sharp fiber optic implanted in ADS to silence the activity of the FOF-ADS projection neurons’ axon terminals (Fig 3A).

First, to test whether this projection has any role in the decision-making process, we silenced the activity of FOF→ADS terminals in either hemisphere during the whole trial (2s inactivation starting 500ms before the stimulus period and ending 500ms after stimulus offset when the animal is free to make their response; Fig 3B,C) on a random subset of trials (25%). This inactivation significantly increased the proportion of choices ipsilateral to the laser side in rats injected with eNpHR3.0 expressing virus (Fig 3C; P < 0.001, nonparametric bootstrap test), with no significant effects on the controls injected with eYFP expressing virus (Fig 3C; P = 0.78, nonparametric boostrap test), suggesting that the projection is involved in the decision process.

Next, we assessed the temporal extent of involvement of this projection, by restricting the inactivation to either early (first half of a 1s long stimulus) or late (second half of a 1s long stimulus) epochs of the evidence accumulation or stimulus period (Fig 3B-C; similar to^16,14^). With this design, it is predicted that if the FOF→ADS projection is involved in conversion of the graded evidence to a categorical choice, then the inactivation should have an effect on behavior only at the end of the stimulus presentation i.e. during the late epoch. Whereas, if the projection is involved in evidence accumulation, a process that occurs throughout the stimulus presentation, inactivations during both early and late epochs should have an effect. Much to our surprise, when we restricted inactivations of the FOF→ADS projection to the early and late epochs, we found that the inactivations led to an ipsilateral bias in both epochs (Fig 3C; early, P = 0.002; late, P = 0.007, nonparametric bootstrap test) with no significant differences in the degree of impairment (P = 0.49) between the two. This is in stark contrast to the effect of whole region FOF inactivations^16^ which caused an ipsilateral bias only during the late epoch. Instead, these results resemble ADS inactivations^14^ which caused an ipsilateral bias in both early and late epochs. Altogether, these results implicate the FOF→ADS projection (i.e. feedback flow under the hypothesis) in the gradual accumulation process, revealing a previously obscured role for FOF neurons. Consistent with this, the involvement of cortico-striatal neurons throughout the stimulus period was also recently reported in mice performing a similar task using somatic inhibition^43^.

Finally, we also examined the effect of inactivations on movements (Supp Fig 5). Terminal silencing slightly but significantly reduced mean movement times for choices both ipsiversive (P = 0.05 Mann-Whitney U test) and contraversive (P = 0.001 Mann-Whitney U test) to the laser (Supp Fig 5A). However, silencing did not increase rates of fixation violations (P = 0.58, nonparametric bootstrap test; Supp Fig 5B). This indicates that the laser did not induce any gross motor impairments affecting animals’ ability to successfully complete trials.

**Figure 5.**
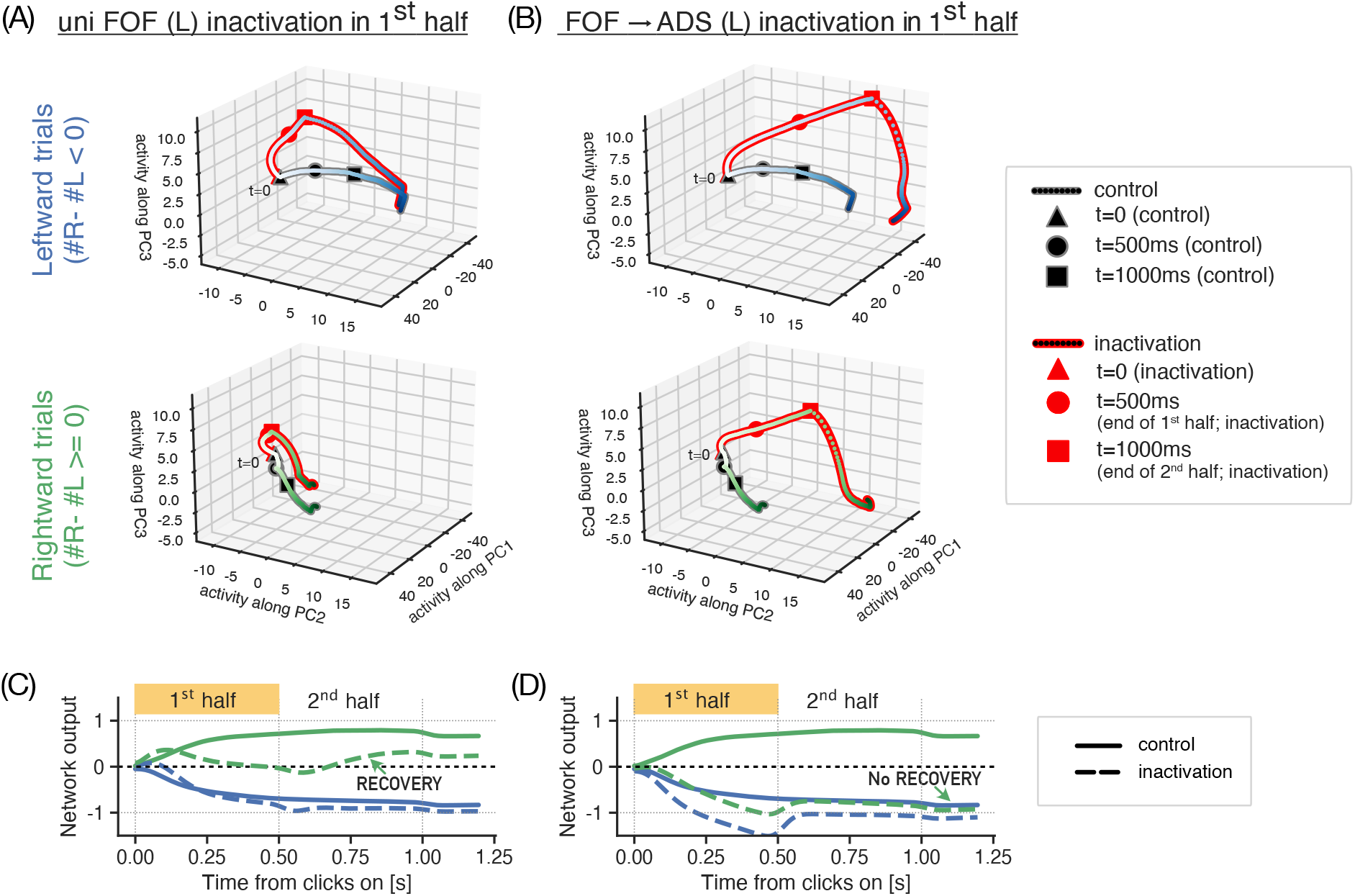
Network activity and outputs during nonselective FOF inactivations reveal recovery dynamics. Low-dimensional projections of network activity trajectories (A,B) and network outputs (C,D) during unilateral (uni; here left hemisphere) inactivations of FOF (A,C) and inactivations of its projections to left ADS (B,D) in the 1st half of the stimulus period. **(A-B)** Neural activity on leftward (top) and rightward (bottom) trials show that inactivation trajectories (lines with red edges, red boxes) diverge from control ones (lines/boxes with grey edges) for both experiments in the first half (until circle markers). However in the second half (from circle to square markers), these recover towards control trajectories during unilateral FOF inactivations (A), but not during FOF→ADS inactivations (B) driving leftward (or ipsilaterally biased) choices. The effect is more prominent in rightward trials (lower panels) **(C-D)** A similar pattern is seen in the network outputs on rightward trials (green lines in C,D), where for both experiments, outputs upon inactivation (dashed) diverge from control (solid) in the first half, recovering and driving correct choices during FOF inactivations (C) but not during FOF→ADS inactivations (D).

### Bilateral perturbations of FOF have effects similar to unilateral FOF

Optogenetic inhibition of FOF inputs to ADS impairs decisions both in early and late stimulus epochs (Fig 3B-C), indicating a role for FOF in the gradual accumulation process, much like ADS. This is consistent with the highly similar neural representations observed in the two regions (Figs 1, 2) and suggests that rather than being incidental, evidence representations in FOF might be important for decision-making behavior. Interestingly, this result also raises the question: why don’t the unilateral whole-region FOF inactivations disrupt decisions during the accumulation period? In mouse M2 interhemispheric recovery was shown to be the reason for lack of impairments observed during transient unilateral inactivations in early task periods^21,44^. So one hypothesis is that the FOFs in the two hemispheres are coupled and can support each other’s recovery when aberrant inputs from the other hemisphere are detected. The localized nature of terminal inhibition (targeting FOF inputs just to ADS, and not to the other FOF) might bypass such a recovery.

We tested this hypothesis by carrying out bilateral optogenetic inactivations in a separate cohort of rats (Fig 3D). We silenced FOF activity by delivering laser bilaterally either during the first half of the stimulus (early inactivation epoch), or the second (late inactivation epoch; same as Fig 3B). We observed a significant decrement in performance only in the late epoch (P < 0.01, nonparametric bootstrap test), while inactivation during the early accumulation epoch did not significantly hamper the performance (P > 0.05, nonparametric bootstrap test). The decrement in performance due to early epoch inactivations was significantly smaller than the decrement in response to late epoch inactivation (P = 0.032, nonparametric bootstrap test). This implies that interhemispheric compensation does not underlie the lack of impairment observed on inactivating FOF early during evidence accumulation. So the question of why projection specific inactivations impair decisions early during accumulation but nonspecific whole region FOF inactivations don’t, remains outstanding.

### A model of distributed implementation of evidence accumulation in corticostriatal circuitry

Our experimental findings are difficult to reconcile with a feedforward organisation of these brain regions, wherein regions have localized functions (as previously hypothesized). They might however be concordant with a circuit that accumulates evidence in a distributed fashion through recurrence and feedback, which allows for adaptive rescue of computations upon perturbation^45,21,46,47,48^. Therefore, we translated our findings into a mechanistic multi-region recurrent neural network (RNN) model that instantiates a recurrent circuit between FOF and ADS, building on the success of RNN models in capturing animal behavior and neural representations during decisionmaking^49,50,51,52,53^. We sought to analyze the behavior of this RNN in unperturbed and perturbed states, so as to identify mechanism(s) through which it can be sensitive to projection specific inactivations but resilient to nonselective whole region FOF inactivations during the accumulation period.

We imposed biological constraints on the RNN, by initializing its weights with possible anatomical connections while following Dale’s law and maintained these constraints on the architecture throughout training (following methods developed in Song 2016^54^). The model’s architecture is schematized in Fig. 4A (also see Supp Fig 6A-B). Briefly, the network included two modules for the two brain hemispheres and each module in turn consisted of three submodules with 50 all-to-all connected neural units each - representing FOF, ADS and the multi-synaptic relay between ADS and FOF (i.e. other basal ganglia nuclei, thalamus, SC, etc.). In the FOF submodules 20% of the neurons were inhibitory whereas in ADS submodules 100% of the neurons were inhibitory, approximating the known distribution of E-I neurons in these brain regions. We modeled long-range projection patterns after empirical observations. The FOF submodule’s projection patterns followed that of cortical long-range pyramidal neurons (IT and PT neurons), ADS projected unilaterally to the relay module and the relay module projected bilaterally to the two FOFs, completing the recurrent loop. The sensory inputs were sent to both FOF and ADS submodules, since it is known that the auditory cortex projects diffusely to both FOF and ADS in the rat brain. Outputs were read out from all neural units in the FOF, ADS and striato-cortical relay submodules.

**Figure 6.**
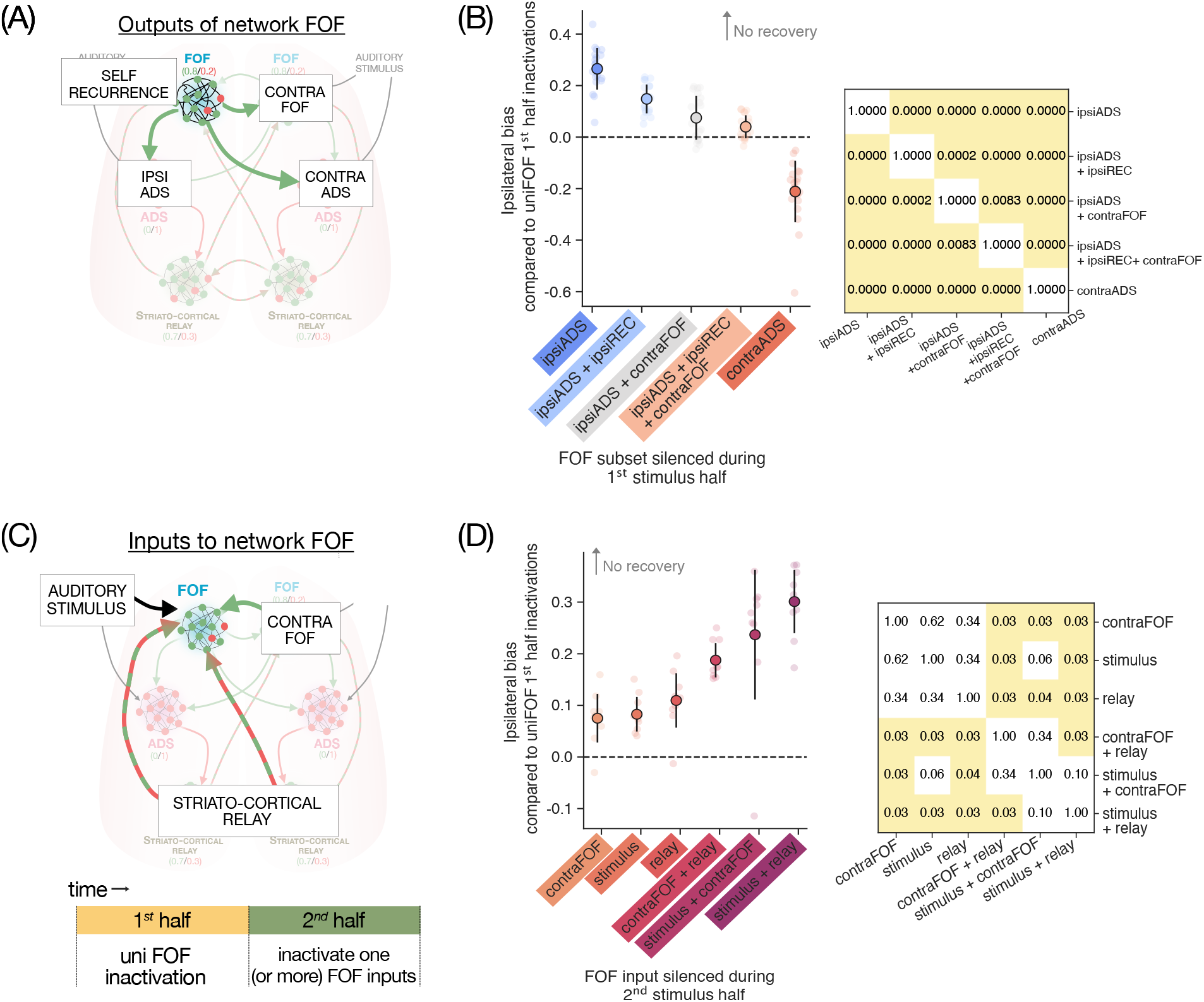
In-silico experiments with RNN to investigate contributions to inactivation dynamics. **(A-B)**: Selectively silencing of different FOF outputs in 1st half invites recovery differentially. **(A)** Schematic of model FOF output projections from a single hemisphere, targeted for in-silico perturbations in the 1st half. Different combinations of output projections were silenced to understand their influence on the inactivation dynamics. **(B)** Summary of in-silico perturbations, showing the ipsilateral bias observed when perturbing a subset of FOF output projections in the 1st half, relative to a baseline of perturbing all its outputs - higher values indicate larger effects and less recovery. Silencing ipsilateral ADS projections has the largest effect and invites the least recovery, additionally silencing recurrent connections and contralateral FOF projections invites partial recovery individually and full recovery in combination. Contralateral ADS projections have no effect to begin with, hence producing less bias than baseline (i.e. negative values). Inset: matrix of p-values for pairwise comparisons between effects, showing that all experiments have significantly different effects. **(C-D)**: Selectively silencing FOF inputs in 2nd half shows multiple contributions to recovery (C) Schematic of model FOF inputs to a single hemisphere, targeted for in-silico perturbations in the 2nd half. (D) Summary of in-silico perturbations, showing the impact of silencing different input projections in the 2nd half on ipsilateral bias, relative to whole FOF silencing in the 1st half. Higher values of bias when a given projection is silenced mean less recovery, implicating that input projection in the recovery process. Inputs from contralateral FOF, auditory stimulus, and striato-cortical relay all have partial influence on recovery, with significantly bigger losses of recovery when they are silenced together. Inset: matrix of p-values for pairwise comparisons between effects, showing that many of these inputs have comparable influence on recovery. P-values were computed with posthoc Wilcoxon test with holm correction for multiple comparisons.

In addition to this multi-region modular structure, our use of the RNN model differs from previous studies in two ways. First, we trained the RNN weights (using backpropagation through time) to perform the evidence accumulation task in a way that matches animal behavior, rather than like an ideal observer (Supp Fig 6B-D). To achieve this, we generated training data by sampling trials from the behavioral model of Brunton 2013^32^. Second, on a quarter of trials during training, we turned down the gain on weights either within an FOF submodule, ADS submodule or FOF inputs to an ADS submodule during either the first or the second half of the stimulus, essentially mimicking our unilateral inactivation experiments (Supp Fig 6E). On these inactivation trials we trained the network to produce an ipsilateral choice, if upon inactivation such a bias was observed empirically. Our use of perturbation data to introduce structure and specialization within a multi-region RNN model is novel, and complements previous approaches that have relied either on differences in neural representations or just the architecture to lend RNNs area-specific identities^55^. We trained the model to successfully reproduce choice behavior on control (Fig 4B) and inactivation trials (Fig 4C). In the model - much like the experimental data - unilateral FOF inactivations introduced an ipsilateral bias only in the second half of the stimulus, but inactivations of FOF to ADS projections and of ADS caused a bias throughout the stimulus. We probed the dynamical structure of these trained models in unperturbed states using the reverse-engineering approach from^56^ and consistent with previous work modeling accumulation of evidence in RNNs^50,57,31^, found approximate line attractors (not shown).

The RNN model’s responses exhibited several features that resembled neural representations, despite not being explicitly trained to do so. The average firing rate of model units, just like neurons from FOF and ADS had heterogeneous temporally patterned responses (Fig 4D). Some neurons encoded the evidence strength transiently whereas other encoded it persistently, capturing the diversity observed in recorded neurons from FOF and ADS. Moreover, the timecourse of stimulus and choice decoding were similar across the model FOF and ADS, as we had observed in our population recordings (Fig. 4E, compare to Fig. 2C and Supp Fig. 1C).

In addition to these similarities in responses, the model was also successful at predicting effects of held-out perturbation. During training, the network was exposed to unilateral inactivations only, so bilateral inactivations of model FOF (Supp Fig 6F) formed a good test of the model’s robustness properties. Prior modeling work comparing the robustness of different architectures^21^ has shown that modular and symmetric networks tend to be robust to unilateral inactivations but not bilateral ones. Therefore, we expected the model to deviate from our experimental findings and show disrupted performance during bilateral inactivations of FOF in the first half of stimulus - unless our novel training protocol with perturbation data was successful at instilling some robustness properties of the real FOF-ADS circuitry into the network. To our surprise, much like the data, the model only showed performance disruptions in the second half of bilateral FOF inactivations, and not in the first half (Fig. 4F, for rat data see Fig. 3D).

### In-silico experiments with the multiregion model reveal network recovery determinants

We examined the responses of neural units during different inactivations by projecting population activity from all submodules onto the first three principal components (Fig 5A, B). Examination of low-dimensional activity trajectories on trials with unilateral inactivation of FOF in the first half revealed that after an initial perturbation, the network activity recovers during the second half (Fig 5A), facilitating normal decision-making behavior. Such recovery was absent during inactivations of FOF projections to an ADS submodule, instead these inactivations sent the population responses along the path that normally drives ipsilateral choices and subsequently gave rise to choices biased towards the inactivated side (Fig 5B). A similar pattern is evident in network outputs, with inactivation trials initially diverging from control trials and subsequently recovering when FOF was inactivated in the first half (Fig 5C), but not when FOF→ADS projections were inactivated (Fig 5D).

In order to investigate which parts of the cortico-striatal circuit could potentially be involved in these recovery dynamics, we took advantage of the precise control we had over RNN activity, and used it as a testbed for performing several *in-silico* perturbations that would be time and cost intensive to run *in-vivo*.

First, we selectively perturbed different output projections from model FOF during the first half individually and in combination, to ascertain what subset of these projections needed to be inactivated to invite the recovery seen in whole-region FOF inactivations. This included self recurrence, projections to contralateral FOF and projections to the ipsilateral and contralateral ADS (Fig 6A, Supp Fig 8A). We found that perturbing projections to the ipsilateral ADS had an inactivation effect in the first half, and invited no recovery in the second half (Supp Fig 8B). In contrast, perturbing projections to contralateral ADS had no inactivation effects at all (Supp Fig 8C). This suggests that the inactivation effects we had observed when perturbing FOF →ADS projections (which were projections to a single hemisphere’s ADS from both FOFs; Supp Fig 6E) may have been largely driven by ipsilateral projections only. Next, we reasoned that inactivating FOF cell bodies should have an effect on the rest of the circuit equivalent to inactivating all of their outputs. We therefore asked, when inactivating the ipsilateral FOF→ADS projection, which FOF output projection inactivations led to the recovery observed when the FOF cells were inactivated. We found that additionally inactivating either of the self-recurrence or contralateral FOF projections led to only partial recovery, while additionally inactivating both of these invited complete recovery (Supp Fig 8D-E). We quantified these effects as the magnitude of the ipsilateral bias caused by these inactivations, relative to the bias caused by unilateral FOF inactivations. By this measure, each additional output inactivation invited significantly more recovery (Fig 6B).

Second, we selectively silenced different inputs to an FOF submodule during the second half, to ascertain which inputs contributed to the recovery in the second half. These included auditory stimulus inputs, inputs from the contralateral FOF and inputs from the striato-cortical relay (Fig 6C). We found that removing any of these inputs individually had minor effects on the ipsilateral bias, only slightly reducing recovery. However, inactivating pairs of these inputs led to significantly more loss of recovery (Fig 6D).

Overall, these experiments suggest that recovery following unilateral FOF perturbations is supported by multiple scales of recurrence in the cortico-striatal circuit - self recurrence within a single hemisphere’s FOF, inter-hemispheric recurrence and striato-cortical recurrence through the relay. Since our in-silico experiments target anatomically valid projections, they make empirically testable predictions about the *in-vivo* cortico-striatal circuitry.

## Discussion

Two separate brain regions in rats, the anterior dorsal stream (ADS) and the frontal orienting fields (FOF), have been respectively linked to the two distinct computations necessary for decision-making based on noisy evidence: the gradual accumulation of evidence and the categorization of that accumulated evidence to reach a final decision^14,16^. However this implied feedforward functional hierarchy going from ADS to FOF has never been directly evaluated. We sought to fill this important gap by simultaneously recording population activity from these two regions during decision-making (Figure 1). Contrary to the feedforward hypothesis, we found that the two regions do not differ in their accumulator encoding, and evidence can be decoded from both regions with comparable accuracy and at comparable timescales. Moreover, we show that this information is shared between the two regions and shows no discernible lead-lag relationship, consistent with a recurrent interaction (Figure 2). Further, we optogenetically silenced FOF axon terminals in ADS (i.e. the “feedback” projection under the hypothesis) during time periods that differentially overlap with the two sub-computations, and found that such a manipulation impairs decision-making during all accumulation periods, unlike nonselective FOF perturbations (Figure 3). Together, these results indicate that the cortico-striatal circuitry participates in evidence accumulation in a distributed manner with ongoing recurrent interactions. We synthesised these findings into a multi-region recurrent neural network model. We successfully trained this RNN with the novel objective of reproducing behavior on control and inactivation trials, and found that it recapitulated neural responses and perturbation effects that it wasn’t trained on (Figure 4). Finally, we investigated the circuit’s response to perturbations by examining the RNN’s activity (Figure 5) and performing in-silico experiments on the RNN, targeting different sets of projections (Figure 6). We found that FOF’s robustness to nonselective perturbations was supported by multiple scales of recurrence in the cortico-striatal circuit.

The similarities we observed in the encoding of task-relevant features in FOF and its down-stream target ADS during decision formation are largely consistent with recent reports in mice in simple decision-making tasks, which found that choice signals in secondary motor cortex (that FOF is a part of) and striatum emerge at indistinguishable time scales^58^ and that striatum reflects summed activity of its cortical inputs^59^. Moreover, our results about encoding of sensory evidence in FOF are in agreement with a recent study of dorsal cortex’s role during decision-making in mice which reported that secondary motor cortex prominently encodes choice independent evidence-related signals^60^. In the random dots motion task^61^, monkey FEF and caudate (the analogues to rat FOF and ADS) have respectively been reported to carry similar accumulation-related signals early during the stimulus presentation with comparable prevalence of trial-difficulty and choice modulated signals during the stimulus epoch (FEF^62^, caudate^36^, FEF and caudate^35,63,64^). This similarity in evidence representation and functional role of cortex (FOF) and striatum has been predicted and is therefore consistent with previous theoretical accounts that have sought to embed the decision-making computation into the cortico-basal ganglia network^3,4^. However similarities with Lo 2006^2^ are unclear, which predicts transient rather than sustained responses in striatum and models striatum’s role as the setting of evidence thresholds.

Despite these similarities in evidence accumulation signals between the two regions, some studies have indeed found differences between FEF and caudate during decision making, particularly in tasks with reward manipulations that require the adaptive tuning of decision thresholds^63,64,65^. Such results fit with wider studies implicating the striatum in action selection^66^ and reward-based habitual learning, and secondary motor cortex in planning goal-directed movements^67,68^. Such distinctions are important avenues for future study and could reveal wider non-overlapping roles of the two regions, but are out of scope for our study that involves a task with fixed, symmetric rewards and is not designed to distinguish between these other functions. Moreover, we have focused our analysis on the encoding and decoding properties of specific task variables in the two regions, such as accumulator tuning, choice and history. We acknowledge the possibility that other views of the neural activity have the potential to reveal subtle differences where we have found none. Indeed work from Luo and Kim^69^ find differences in the geometry of dynamics in the two regions. Whether such distinctions influence the functional roles of these regions is an open question, and one that we leave for future work to investigate.

To understand if the evidence representations in FOF and ADS are related, we used reduced rank regression to identify dimensions in the neural population space that captured co-fluctuation in the trial-to-trial activity of FOF and ADS over and above those introduced by external task events and defined a functional communication subspace^70,26,71,72,73,41^. We found that during the stimulus period, the communication subspace carries stimulus-related information such that evidence can be decoded fairly well from just the activity present in the communication subspace. This did not have to be the case, as using similar approaches recent work has identified patterns of activity or information that are shared between brain areas as well as those that are kept private within an area. For instance, it has been reported that majority of the variance in V1 is kept private and that V1 only selectively shares information related to a few of its activity patterns with V2^20^. In rat M1 and M2 it has been shown that shared dynamics across areas identified using a related method (canconical correlation analysis; CCA) become more related to task performance with learning^74^. Similarly in mice, CCA was recently used to show that the content of communication between A1 and mPFC changes during different contexts based on a control signal from mPFC, this helps select revelant sensory information during contextual decision making^75^. Such communication subspaces defined based on noise correlations do not distinguish between correlations due to direct communication from those induced by communication arriving from common inputs to the two regions. While our projection-specific inactivations support a causal role for the interaction from FOF to ADS, our conclusions about communication of evidence between the two regions can be further strengthened if decrements in evidence-related information are observed upon perturbation of one of the areas, in paired perturbation and recording experiments.

Through specifically targeting the cortico-striatal pathway and silencing FOF axon terminals in ADS, we revealed a role for FOF throughout the gradual accumulation process. This finding might seem at odds with the past work that had concluded using unilateral silencing of FOF somas, that FOF’s causal role is restricted to the decision commitment period and does not extend to the accumulation period^16^ – however it is in line with recent exciting work highlighting how different projection pathways from the same cortical region can carry different information and differentially impair behavior upon perturbation^22,76,77,78,79,80,43,81,82,83,84^ (further discussion^66,85^) producing effects that could be otherwise obscured by whole region perturbations. A notable example of this comes from an auditory frequency discrimination task in rats^76^, where perturbations of auditory cortex neurons with high (low) frequency tuning did not bias animals’ choices towards high (low) responses, but selective activation of striatum projecting auditory cortex neurons with high (low) frequency tuning biased the animals’ choices as predicted. Also of note is a recent study in mice^43^, which has also proposed a role for corticostriatal projections from frontal cortex throughout gradual accumulation of evidence. Our results add an important piece of evidence in favor of the emerging principle that projection-specific perturbations can unmask effects not visible in whole-region perturbations^22,76,24^. Given recent controversies about the causal roles of different regions in evidence accumulation^86,15,25,87,88^, our approach might offer a way to further clarify the roles of these brain regions and help with identifying the key communication channels operant during decision-making.

While the encoding/decoding properties of units in our network resemble those of the empirical data, a complementary approach would be to train the RNN such that individual units in the network additionally match observed neural firing rates^89,90,91,92,93^. It remains an exciting empirical question of how these different training objectives, algorithms, and constraints alter the dynamics of the network and which objectives make better predictions. Past studies seeking to model the complex web of multi-region interactions that give rise to cognition^94,95,96,47,97,48^ (reviewed in^98,55^) by training RNNs have relied either on differences in neural representations or just the architecture to lend RNNs area-specific identities. We believe our use of projection-specific perturbation data to introduce structure is novel and is likely to tighten the correspondence between the models and data (also see^99^). We are hopeful that this approach of using multi-region RNNs in tandem with projection-specific perturbations and simultaneous recording will offer a powerful framework for generating hypotheses and intuitions about how recurrently connected brain regions perform decision-making in a distributed fashion.

## Methods

### Subjects and housing

A total of 19 male Long-evans rats (*Rattus norvegicus* from Hilltop, PA) were used for this study. Of these, 5 animals were used for neural recordings and 14 for optogenetic experiments. Investigators were not blinded to experimental groups during data collection or analyses. Animal use procedures were approved by the Princeton University Institutional Animal Care and Use Committee (IACUC #1853) and were carried out in accordance with National Institute of Health standards.

Rats were housed in Techniplast cages and all training and testing were conducted during the dark cycle. Animals that trained during the day were housed in a 12 hr reverse light cycle room. Animals were pair housed whenever possible, but were always single housed after optic fiber or Neuropixels implantation to prevent damage to the implant. Rats had free access to food but were placed on a controlled water schedule in order to motivate them to participate in behavioral task for water rewards. They obtained water rewards during behavioral training sessions (1-5hrs). Following behavioral training rats received either an unlimited water supplement in an ad lib access period of up to 1hr or a controlled supplement so that their total access to water within a period of 24hrs equalled 3% of their body weight.

### Neuronal recordings and analysis

#### Acquisition and pre-processing

We followed the general surgery and Neuropixels probe implantation methods for recording chronically in freely moving animals from^34^ to record unihemispherically from FOF and ADS of 5 rats. The Neuropixels probes were assembled in a compact 3D printed casing, printed in-house using Formlabs SLA printer. This assembly allowed the probe to be stereotaxically manipulated, provided electromagnetic shielding, prevented it from contacting biological fluids or other adhesive materials applied during surgery and imparted mechanical protection to the implant for robust chronic tethered recordings. We targeted FOF and ADS with one penetration, by implanting the probe at AP 1.9mm and ML ±1.15mm from bregma, at an angle of 15^°^ in the coronal plane. Before implantation, the probe shank was dipped for several seconds in a dye (DiI, 1-2 mg/ml in isopropyl alcohol) to allow for histological reconstruction of probe location. We inserted at least 6mm of the probe shank into the brain; the actual depth of implantation varied between animals.

Electrophysiological recordings were made using either commercially available Neuropixels 1.0 acquisition hardware^100^ or the earliest test-phase IMEC acquisition hardware. We used SpikeGLX software to acquire the data. On every session we recorded from 384 sites arranged in a checkerboard pattern spanning banks 0 and 1 of the probe. The data were automatically spike sorted offline using Kilosort2 with default parameters and then manually created using JR-Clust’s GUI^33^. During manual curation, each unit detected by Kilosort2 was inspected, if the events comprising the unit had near zero amplitudes or non-physiological waveforms then the entire unit was discarded. If the unit was judged to be comprised of multiple distinct waveform then it was split into two or more units and finally spatially neighboring units were compared using cross-correlograms, drift patterns and waveform similarity to determine if they should be merged. After following these steps, all units with < 5% refractory violations were considered as reflecting spikes from a single neuron.

#### Analysis of neural recordings

For all analyses, we analyzed neurons that had an average firing rate greater than 1Hz during the stimulus period, neural responses were aligned to stimulus onset and included only up to the time of movement onset for each trial. To compute the average population response to different stimulus strengths, we selected neurons that had large differences in their firing rates during the stimulus period for trials that subsequently resulted in rightward versus leftward choices (|auc −0.5| > 0.05, *P* < 0.05, Receiver operating characteristic). The choice that produced the larger response was defined as the preferred side. For each neuron, firing rates for individual trials were calculated by smoothing the spike trains with a causal half-Gaussian filter with 0.05s standard deviation, and then normalizing by neuron’s mean response at stimulus onset.

##### Neural tuning curves

Following Hanks 2015^16^, we fit the trial-by-trial behavioral model developed in^32^ separately to each rats’ choices. In order to best describe the behavior on recording sessions and account for nonstationarities, we first fit the model to choices from sessions (min 30, upto 90) prior to recording. We then used the best-fit parameters and their estimated confidence intervals to impose Gaussian priors on the parameters and refit the model to behavior from just the recording days. The mean of the Gaussian prior was set to the maximum likelihood estimate of the parameters, and the standard deviation was set to 2 times the estimated confidence interval. Once fit, we obtained the trial-by-trial, moment-by-moment estimate of the likely accumulator variable trajectories while taking into account the choice made by the rat from the model. This was computed by selectively propagating accumulator values that are consistent with rat’s choice - at the end of the stimulus - backwards in time.

We collated across trials and used these estimates of the accumulator value to compute the joint probability distribution between accumulator value, time and observed firing rates for each neuron. Following past work, the neural time was assumed to lag behind the time in the model by 100ms. Our results are invariant to small deviations (50 − 100ms) from this assumed lag. Firing rates were computed by smoothing the spike trains with a Gaussian filter with 0.75s standard deviation. We marginalized the joint probability distribution over time to extract the response of the neuron given the value of the accumulator. We analysed the time period between 0.15 and 0.5s from stimulus onset and responses of side-selective neurons (|auc −0.5| > 0.075, *P* < 0.05). These settings yielded firing rate maps that were largely stable in time for different accumulator values. We accounted for differences in the dynamic ranges of the neural response, by scaling the responses to span the range from 0 to 1. We then fit the estimated relationship between scaled neural responses *r* and the accumulator value *a* with a four-parameter sigmoid:

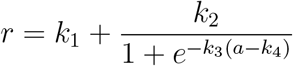

At zero-crossing, the slope of this equation is given by *k*_2_*k*_3_*/*4. We compared this slope parameter for individual neurons across the two regions and used the Mann-Whitney U test to determine whether the medians of each region’s population distribution were significantly different.

##### Decoding analyses

We trained linear decoders to predict the cumulative difference in number of right and left clicks (Δclicks) from neural population activity. To be able to compare decoding performance from the two regions, we controlled for differences in the number of recorded neurons by sub-sampling neurons from the region with higher number, while maintaining the overall distribution of side selectivity. We considered time points aligned to stimulus onset, and assumed a neural encoding lag of 100ms. Δclicks during the trial and neural responses were both binned into 50ms bins, neural responses were further smoothed with a Gaussian kernel with 75ms standard deviation. Neural responses were then assembled into a design matrix *X* with dimensions *T* × *N*, where *T* is equal to the total number of time points (time points from stimulus onset to termination × number of trials), and *N* is the number of neurons. Similarly, Δclicks were concatenated across trials to yield *Y* with dimensions *T* × 1. We fit one set of regression weights *β*_*st*_ for all time points such that *Y* = *Xβ*_*st*_ and imposed L1 penalty. We performed 10-fold cross-validation to assess the decoder’s performance. Within each fold the strength of regularization was estimated independently using 10-fold cross-validation. Fitting was performed using the sklearn package in Python^101^. To determine if the decoding performance or its timecourse was different between the two regions, we performed a two-way repeated measures ANOVA with time (time bins from stimulus onset) and area (FOF, ADS) as factors using Pingouin package in Python^102^.

To decode binary variables, choice and past trial’s choice we used logistic decoders. We assembled the design matrix similarly, however without any neural lags since the target variables are constant for a trial. We duplicated the eventual or past choice for all time bins from a trial to construct the *Y* variable. We fit the decoders with L2 penalty, once again with 10-fold cross-validation. Again, we used sklearn package in Python for fitting^101^. While estimating the decision variables or DVs using the decoders trained to predict choice, we binned the neural activity into 1ms bins and convolved them with a Gaussian kernel with 25ms standard deviation before projecting onto the decision hyperplane. These parameters were picked based on recovery analysis on simulated synthetic datasets. However, our results are not sensitive to small variations in these parameters.

##### Reduced rank Genralized linear model (RR-GLM)

We used RR-GLM to define the FOF → ADS and ADS→FOF communication subspaces. For this we fit a GLM with a softplus link function to the entire population activity. The model was optimized to estimate the firing rate of neurons at each time point *t*, assuming Poisson emissions and given a set of time-varying task parameters. Neural spikes during a period of 2s from initiation of a trial were fit. The following task variables were included: trial start, stimulus start, left and right clicks and current trial’s choice (separate variables for the left and right choice were included). The task variables were linearly convolved with a kernel to allow for time-varying affect on neural spiking. Kernels for choice were allowed to be acausal, whereas all other kernels were causal in their timing. Following past work^103,104^, to enforce smoothness onto these temporal kernels, we parameterized each kernel using raised cosine bases. Hyper-parameters, such as number and duration of these kernels, were optimized through cross-validation on a small grid. Additionally, temporal kernels with raised cosine bases were fit for the simultaneously recorded population from the other region, incorporating a one-time step delay between the regressor and regressand populations. The regressor activity was first projected onto a lower rank subspace to which the convolutional kernels were applied. We binned the neural spiking activity into 5ms bins. Both the linear weights for the kernels and the dimensions spanning the lower-rank subspace were fit simultaneously by minimizing negative Poisson log-likelihood of the spikes with an L2 penalty, using the Adam optimizer^105^ in Pytorch^106^. The regularization strength was chosen using cross-validation. For decoding from the communication subspace, procedures identical to full population space (section above) were followed.

### Optogenetic perturbations and analysis

#### Optical fiber chemical sharpening

We followed methods similar to Hanks 2015^16^ and used standard off the shelf 50-125*μm* FC-FC duplex fiber optic cables (FiberCables.com). The metal casing of the connector and the outer protective layers of the cable were removed, yielding a 1.5cm of fiber optic cable with inner plastic coating (typically clear) intact. To etch the fiber tip, 2-2.5mm of the fiber tip was submerged in a 5ml Eppendorf tube containing 48% hydrofluoric acid, topped with mineral oil. Over the course of 17mins, the fiber tip was slowly pulled out of the acid using a motor (Narishige) producing a taper in the inner plastic coating. Then the speed of the motor was increased and the motor was run until the tip was entirely removed from the acid (typically 13mins), producing a sharp fiber tip, usually with uniform and broad light scatter. We measured the laser beam transmission efficiency of the etched fibers using an integrating sphere photodiode power sensor (Thor Labs). Fibers that did not produce sufficiently broad or uniform scatter or had efficiency < 85% were discarded.

#### Optogenetic virus injection and fiber implantation

We followed previously described procedures for general surgery and virus injections^107,16^; here we describe procedures specific to this study. We bilaterally injected adeno-associated virus, either AAV5-CaMKIIa-eYFP-eNpHR3.0 or AAV2/5-CamMKIIa-EYFP-WPRE-hGHpA using a Nanoject (Drummond Scientific) in FOF. Five closely spaced injection tracts were made in each FOF’s craniotomy. The tracts were typically arranged 500*μm* apart at the 4 vertices and the center of a cross spanning AP, ML axes, centered at AP 2mm ML ±1.3mm. In each tract, 16 injections of 14.1*nL* were made every 100*μm* in depth, starting at 200*μm* below brain surface i.e. from 0.2-1.8mm. Virus was expelled at 20*nL/s* and the injections were made once every 10*s*. At the final injection in a tract the pipette was left in place for at least 2min before removal. In each craniotomy a total of 1.128*μl* of virus was injected over a 30min period consisting of 80 separate injections.

For bilateral FOF silencing, chemically sharpened optical fibers were then lowered down the central injection tracts to a depth of 1mm. For inhibition of FOF terminals in ADS we made additional craniotomies centered over ADS (AP 1.9mm, ML ±2.4mm) and lowered etched fibers to a depth of 3.5-3.9mm. The craniotomies were filled with kwik-sil (World Precision Instruments) and the fibers were secured to the skull with viscous dental composite (Absolute Dentin, Parkell). Optical fibers were then enclosed in a custom 3D printed casing and the implant was filled with dental acrylic, allowing only the FC connectors to protrude. Before behavioral experiments, viral constructs were allowed to develop for 6-8 weeks for FOF silencing, and for 10-12 weeks for terminal silencing. Accurate viral injection targeting and expression was verified histologically.

#### Optogenetic perturbation

For optogenetically perturbing neural activity during behavior, a fiber rotary joint (Princetel) was mounted on the ceiling of the behavioral chamber. In the beginning of a session, the FC connectors in the rat’s implant were connected to the patch cables attached to the fiber rotary joint. The rotary joint was connected to the laser beam source. For FOF inactivations, the laser beam from a 200mW, 532nm laser (OEM Laser Systems) was used. To deliver illumination bilaterally, the laser beam was split into two beams of roughly equal power (∼ 25mW) using a beam splitter (Doric). An OBIS 594nm (Coherent) laser beam of power 25-33mW was used for silencing of FOF axon terminals in either left or right ADS (unilateral silencing). Illumination was delivered continuously on a subset of trials (25%) by opening and closing a shutter with a 5V TTL pulse.

In the inactivated trials, light was delivered during one of three different epochs. The first epoch was 2000ms long, starting on trial initiation and ending 500ms after the termination of the stimulus. We call this “whole-trial” inactivation and this type of stimulation was delivered to a cohort of 6 rats (10 hemispheres). The second and third epochs were 500ms long and spanned the first and second halves respectively of a 1s long auditory stimulus. We refer to them as “early” and “late” inactivation epochs respectively. Early and late inactivations were done in sessions separate from the whole trial inactivations in 5 rats (8 hemispheres; 3 of these rats also belonged to whole-trial inactivation cohort). In sessions with early and late inactivations, 50% of the trials had 1s long stimulus. Duration of the stimulus on rest of the trials was uniformly distributed between 0.2-1s. On half of these 1s long stimulus trials, light was delivered either during early or late epoch, and the other half served as control trials.

We measured behavioral bias resulting from optogenetic inactivation by following methods used in^16^ and^14^. First, we binned the trials on the basis of stimulus strength and for each of the 10 binned stimulus strengths, we computed the percentage of trials during which the rats chose the side ipsilateral to the optically stimulated side on control and inactivation trials. The mean difference across stimulus strengths in this ‘go ipsi’ rate between inactivation and control trials measured the bias caused by optogenetic inactivation. A positive bias represents an increase in ipsilateral choices upon laser stimulation. Confidence intervals and statistical comparisons for this bias parameter were calculated using nonparametric bootstrap procedures.

### Multi-region recurrent neural network model

We trained an RNN with biological constraints on connectivity to perform the evidence accumulation task from^32^. The RNN is composed of standard firing-rate units (*N* = 300) with a rectifying nonlinearity and *τ* = 30*ms* as the time constant of decay of network units. RNN dynamics are governed by the equations:

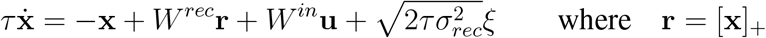

where the variable **x**(*t*) is an *N* dimensional vector containing the activation of neural units in the network. The activations are mapped to the corresponding firing rates **r**(*t*) by passing **x** through the threshold linear nonlinearity which maps individual activations to positive firing r ates: *x* _*i*_ if *x*_*i*_ > 0 and 0 otherwise for *i* = 1, …, *N*. Recurrent connection weights in the network are given by the *N* × *N* matrix *W* ^*rec*^ and the connection weights from the inputs to network units are given by the *N* × *N*^*in*^ matrix *W* ^*in*^. The network receives 2-dimensional (*N*^*in*^ = 2) time-varying inputs, **u**(*t*) = [*s*_*L*_(*t*) *s*_*R*_(*t*)]^T^, which represent the left clicks and right clicks respectively (Supp Fig 6 B-D). Noise intrinsic to the network is represented by *N* × 1 dimensional *ξ*(*t*), independent white noise with zero mean and unit variance. The network output *z*(*t*) is read out linearly, as a weighted sum of the firing rates of network units, with weights *W* ^*out*^ and a bias *b*:

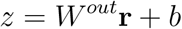

#### Network inputs, perturbations and targets

The network received 2 inputs **u**(*t*) = [*s*_*L*_(*t*) *s*_*R*_(*t*)]^T^ at every time step. The two inputs correspond to the randomly timed discrete auditory evidence that is presented to the rats during the task, from speakers to their left and right sides respectively. These inputs are zero except when a left/right click occurrs. The click times were sampled from Poisson processes, with Poisson rates for left/right inputs sampled independently on each trial, from the same distribution as the one used for rats. At the times when left clicks occurred the average input was equal to −1, however we added independent Gaussian noise *η* with zero mean and variance 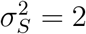 to each click, hence the input for any given left click was −(1 + *η*). 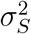 represents the variance of sensory noise accompanying each sensory input, and was set to match the average sensory noise variance measured in rats performing this task^32^. Similarly, when a right click occurred, the magnitude of input *s*_*R*_ was set to 1 + *η*. The overall timing of these stimulus inputs mirrored that of the real task. On each trial, the stimulus duration i.e. the time during which clicks played (in s) was sampled uniformly from the interval [0.2, 1.0] and was preceded by a variable delay such that the stimulus always ended at 1.5s from the trial start.

Choice targets for each trial were obtained by passing stimulus, history and lapsing inputs to the behavioral model from Brunton 2013^32^. If the target choice was towards right (left), then the target was set to +1 (−1) in the 200ms following termination of stimulus, i.e. from 1.5 to 1.7s. On a quarter of trials, we turned down the gain of certain recurrent weights in *W*^*rec*^ by a factor of 0.1 so as to simulate unilateral inactivations of FOF, ADS or FOF inputs to ADS during either the first or second half of a 1s long stimulus. On these trials, if an ipsilateral bias was observed experimentally then the choice targets were not obtained from the model but instead always set to the ipsilateral side.

#### Network connectivity and training

We engineered the recurrent weight matrix *W*^*rec*^ so as to capture the major anatomical features of the FOF-ADS circuitry. The network included two modules for the two brain hemispheres (150 units each) and each module in turn consisted of three submodules with 50 all-to-all connected neural units - representing FOF, ADS and the multi-synaptic relay between ADS and FOF (i.e. other basal ganglia nuclei, thalamus, SC, etc.). All units in the network followed Dale’s law. In other words, all recurrent projections from a unit were either positive or negative but never both. We refer to the units which send out positive recurrent projections as excitatory units and the ones with negative projections as inhibitory units. Within each submodule, we set the ratio of excitatory to inhibitory units to match the known distribution of E-I neurons in these brain regions. Therefore, FOF submodules had 20% inhibitory units and ADS had 100%. For the relay submodule we set the ratio to 30%.

In FOF submodules, inhibitory units projected locally within the submodule, and excitatory units projected to other submodules in a manner aiming to mimic the long-range projections of cortical pyramidal neurons^108^. About 30% of the excitatory neurons from an FOF submodule followed the pattern of projections observed in cortical IT neurons - they sent projections to the other FOF submodule and the two ADS submodules. Another 30% followed projection patters of cortical PT neurons - they sent projects to the ipsilateral ADS submodule. The remaining FOF excitatory units only had local recurrent connections. The ADS submodules projected ipsilaterally to the relay module, since we are not aware of any cross-hemipshere projections between the two hemispheres. We do not differentiate units in ADS as belonging to D1/D2 subtypes etc. Given the abstract nature of the relay submodule, we put minimal constraints on its recurrent connections and it was allowed to project bilaterally to the two FOFs and to the other relay submodule with both excitatory and inhibitory projections.

To initialize *W*^*rec*^, we first sampled a matrix of values from a normal distribution with zero mean and variance 1*/N* while enforcing the connectivity patterns as detailed above. Then we balanced the average weights of excitatory and inhibitory inputs into each unit and set the spectral radius to 1.3.

All units in the FOF and ADS submodules received the inputs i.e. *W*^*in*^ projected only to FOF and ADS units, and outputs were read out from all units in the network. Input and outputs weight were initialized using samples from Glorot normal distributions^109^.

While training, to enforce the Dale constraints on connectivity we parameterized the recurrent weight matrix *W*^*rec*^ as a product of a non-negative matrix *W*^*rec*,+^ and a diagonal matrix *D* of 1s and −1s identifying excitatory and inhibitory units i.e. *W*^*rec*^ = *W*^*rec*,+^*D*. We maintained the sparse connectivity across submodules by further performing element-wise multiplication with a connectivity mask *M*^*rec*^, such that *W*^*rec*^ = *M*^*rec*^ ⊙ (*W*^*rec*,+^*D*). We simulated the network’s dynamics using Euler updates with Δ*t* = 5ms and generated 240, 000 trials for this, with minibatches of 64 trials each. We optimized *W*^*in*^, *W*^*rec*^, *W*^*out*^, *b* and *x*(0) to minimize the mean-squared error (MSE) between the network output and targets, using Adam^105^ in TensorFlow^110^. The methods used for analysis of activity and perturbation response of the multi-region models were identical to that used for rat data.

## Acknowledgements

We thank members of the Brody lab for experimental support and helpful feedback throughout the project. We also thank Jovanna Teran and Brody lab technicians for assistance with rat training. We are grateful to Jonathan Pillow, Ilana Witten, Joshua Gold, David Zoltowski and Aditi Jha for their suggestions and discussions. This work was supported by NIH grant R01MH108358 awarded to C.D.B as well as a grant from the Simons Foundation (Grant number: NC-GB-CULM-00003118-03) awarded to C.D.B.

## Author Contributions

D.G., C.D.K. and A.G.B. collected data. T.Z.L, A.G.B. and V.E. helped with data collection and pre-processing. D.G. did the analysis and modeling, and wrote the initial draft of the manuscript. All authors provided feedback on the manuscript. C.D.B. oversaw all aspects of the project.

## Supplementary materials

**Supplementary Figure 1.**
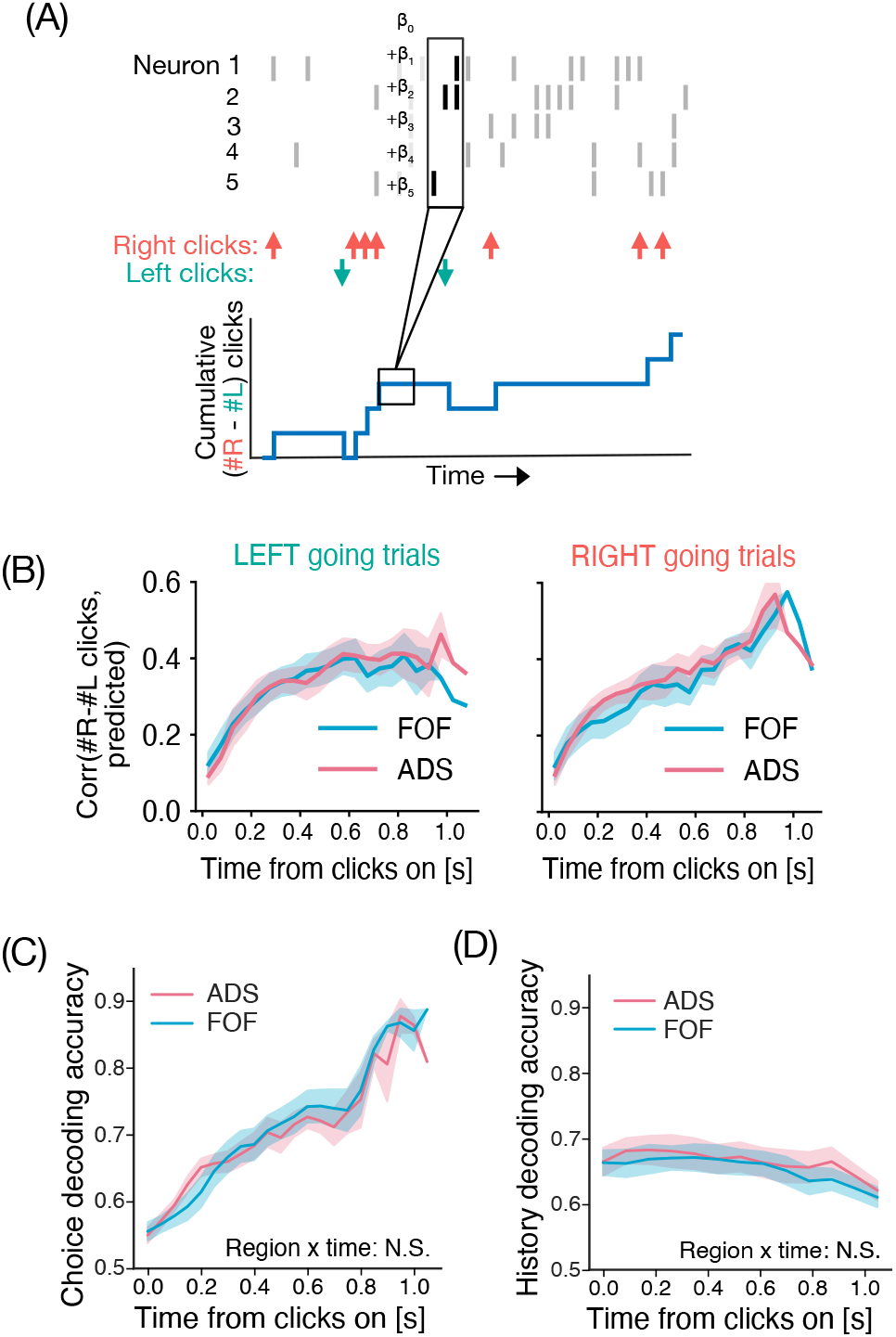
Similar strength and timecourse of stimulus, choice and history decoding in FOF and ADS. **(A)** Schematic showing the linear decoder designed to predict the cumulative click difference in the number of right and left clicks at any given time during a trial using a linear combination of neural activity from FOF or ADS. We assumed a neural encoding lag of 100ms and regularized the linear weights on neurons with L1 penalty using cross-validation. **(B)** Stimulus decoding performance (mean ± sem) of the linear decoder on held-out time points as a function of time from stimulus onset (n = 12 sessions) performed separately for left (left panel) and right (right panel) going trials to control for choice. The two areas show high decoding performance which evolves with a similar timecourse (P> 0.05, two-way RM ANOVA). **(C)** Cross-validated choice decoding accuracy as a function of time from stimulus onset for FOF (aqua blue) and ADS (pink) neural activity (n = 12 sessions). Time courses and strength of choice decoding from the two populations did not differ significantly (P> 0.05 two-way RM ANOVA). Decoding was performed with an L2 regularized logistic decoder, while controlling for number of neurons from the two areas. **(D)** Same as C but decoding for choice from past trial (n = 12 sessions). No significant differences were observed between the two populations (P > 0.05, two-way RM ANOVA)

**Supplementary Figure 2.**
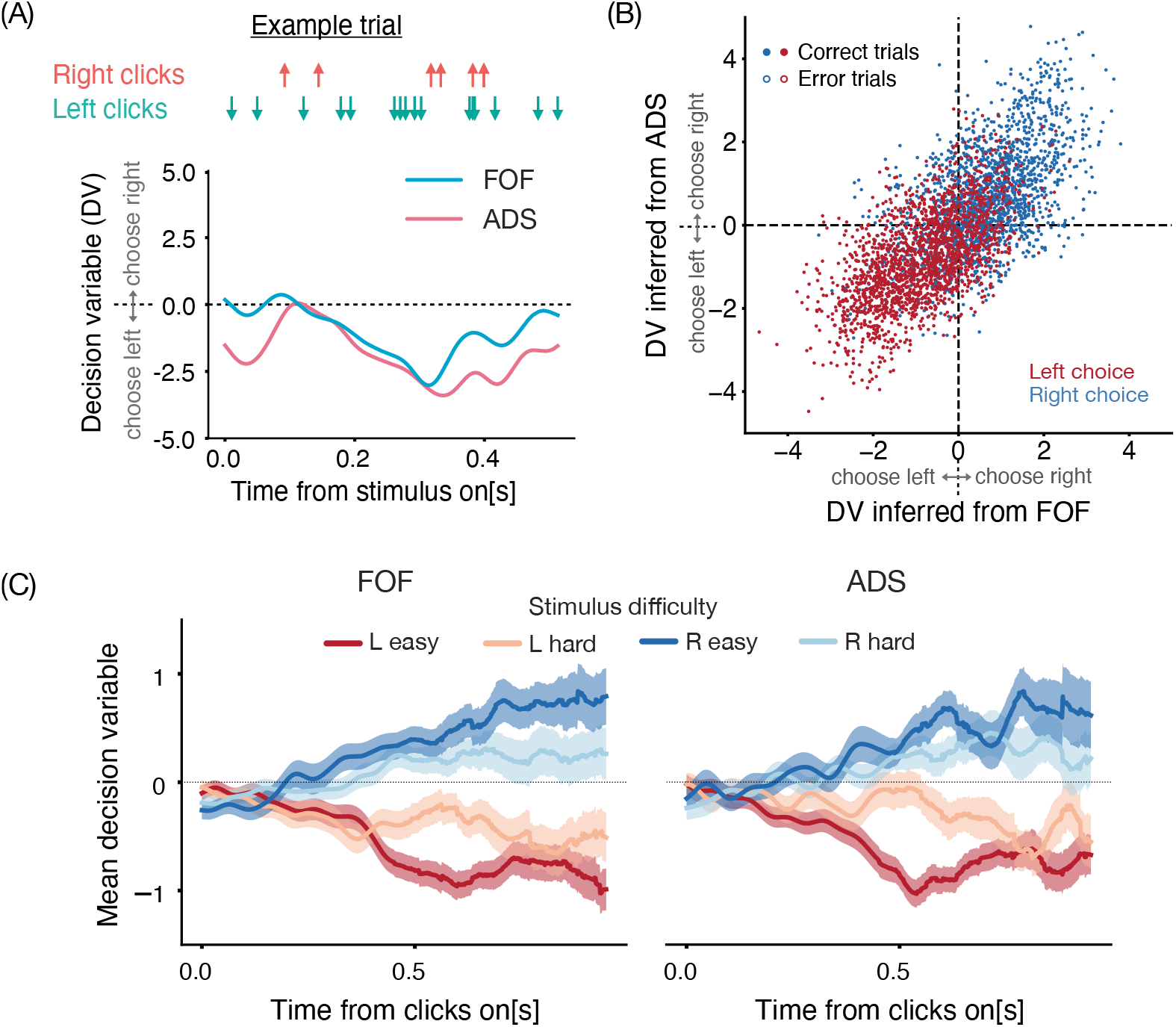
Decoding decision variable from FOF and ADS population activity. **(A)** Example trial showing the trajectory of the decision variable (DV) decoded from the simultaneous population activity of FOF and ADS (blue, pink line respectively), using the logistic decoder (see Methods). The DV represents the log odds ratio of making one choice over another, with higher positive (negative) values indicating a stronger likelihood of rightward (leftward) choices. Both regions have comparable DV time-courses, and seem to be influenced by the sequence of right and left clicks (coral, green arrows respectively). **(B)** Scatter plot comparing DV values inferred from FOF (x-axis) and ADS (y-axis) activity at the same time-points from an example session. Dots represent time-points across trials, with filled (unfilled) circles indicating correct (incorrect) trials. Points are color-coded according to the actual choice of the animal, with blue (red) indicating rightward (leftward) choices. The DV inferred from both areas shows good correspondence with each other and the true choice. **(C)** Mean DV trajectories from FOF (left) and ADS (right) over time from an example session and separated by stimulus difficulty (red - easy leftward stimulus, light red - hard leftward, blue - easy rightward, light blue - hard rightward). Shaded regions indicate S.E.M. Both regions show similar DV time-courses and similar dependence on stimulus difficulty.

**Supplementary Figure 3.**
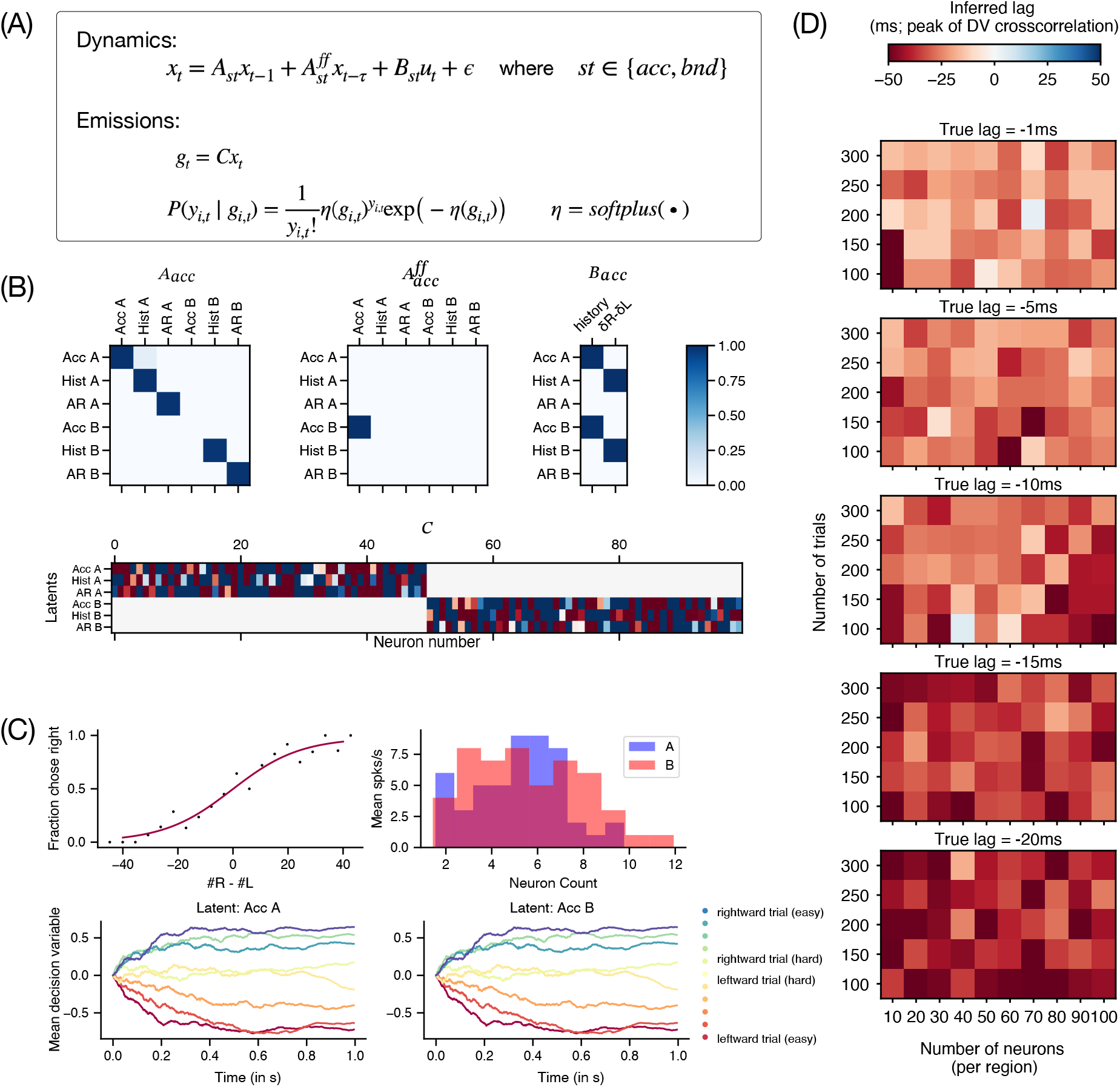
Simulations used to validate the decision variable cross-correlation analysis. **(A)** Equations used to simulate two regions with feedforward flow of accumulator dynamics. (Top) The low-dimensional latents in the two regions *x*_*t*_ represent accumulator variables (Acc), history (Hist) and autoregressive (AR) terms that evolve according to linear dynamics with dynamics *A*_*st*_ that are different during accumulation (*st* = *acc*) and bound-hitting (*st* = *bnd*). During accumulation, one region’s accumulator dynamics (region B) are influenced by feed-forward inputs from the other region’s accumulator (region A) according to 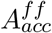 with delay *τ*. Both regions receive external inputs *u*_*t*_ (including stimulus clicks and history) according to *B*_*st*_ and Gaussian noise *ϵ*. (Bottom) The mean firing rates *g*_*t*_ of neurons in the two regions reflect a high-dimensional projection *C* of the aforementioned latents *x*_*t*_. These are then transformed through a softplus function (to enforce positive firing rates) into Poisson distributed spikes *y*_*t*_. **(B)** (Top) Schematized *A*_*acc*,_ 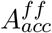 and *B*_*a*_ *cc* (Bottom) Linear weights *C* that transform latents (y-axis) into mean firing rates of individual neurons (x-axis). **(C)** (Top left) Fraction chose right based on thresholded values of the accumulator, showing a monotonic dependence on the stimulus. (Top right) Firing rate distributions in the two regions showing substantial heterogeneity and overlap. (Bottom) Mean decision variables across trials of the same stimulus difficulty for the two accumulator latents, showing a separation by difficulty. **(D)** (Top to bottom) Inferred lag measured by the peak of the decision-variable cross-correlation, for increasing true lags between the regions as a function of number of neurons (x-axis) and trials (y-axis). Even though the exact value of lag is not recovered for low trial and/or neuron counts, the direction of flow can be accurately inferred. Increasing color intensities show that the measured peak lag is sensitive to the true lag values.

**Supplementary Figure 4.**
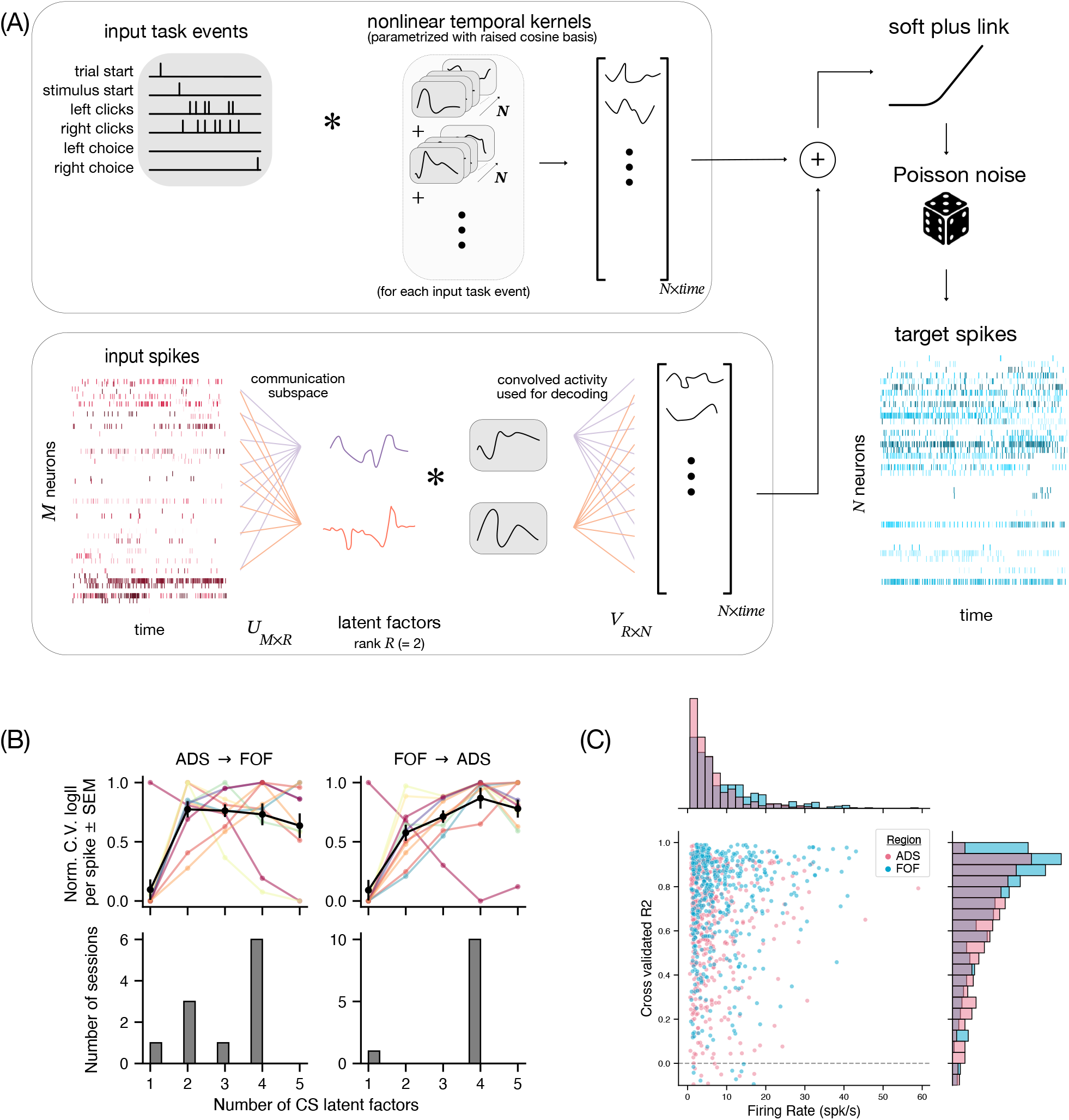
Reduced rank GLM captures the low-dimensional communication subspace between FOF and ADS. **(A)** Schematic of the reduced rank GLM approach. The target variables are sequences of spikes for each of N neurons in the output region (blue rasters), which are modelled as a generalized linear model with a soft plus link function and poisson noise distribution. The predictors consist of task events (top box, left panel) and sequences of spikes from M neurons in the input region (bottom box, red rasters). Task events are convolved with nonlinear temporal kernels parameterized with raised cosine bases - one per task event per output neuron - allowing them to have temporally extended effects (top box, grey boxes). Spikes from the M neurons in input region are first projected into a reduced R-dimensional communication subspace using learnt weights *U*_*M*×*R*_, yielding a small number of latent factors (bottom box, colored curves). These latent factors are then convolved with temporal kernels (grey boxes) and influence the N neurons in the output region through learnt output weights *V*_*R*×*N*_. **(B)** Required dimensionality of the communication subspace. (Top) Normalised cross-validated log likelihood per spike, as a function of the dimensionality of the communication subspace. Colored curves individual sessions, black represents mean across sessions. (Bottom) Distribution of the optimal dimensionality of the communication subspace across sessions. Based on both these measures, the smallest dimension required (≤ 4) was used for each session, with most sessions warranting a 4-dimensional communication subspace. **(C)** Cross validated r-squared values across sessions of the reduced rank GLM (ADS = 0.64 ± 0.01, FOF = 0.73 ± 0.01 mean ± sem), plotted as a function of the session average firing rates - colors represent different directions of communication, with ADS (pink) or FOF (blue) as output regions. In ADS (FOF), 0.12 (0.09) fraction of cells had R2 < 0. These cells were excluded while computing the mean R2.

**Supplementary Figure 5.**
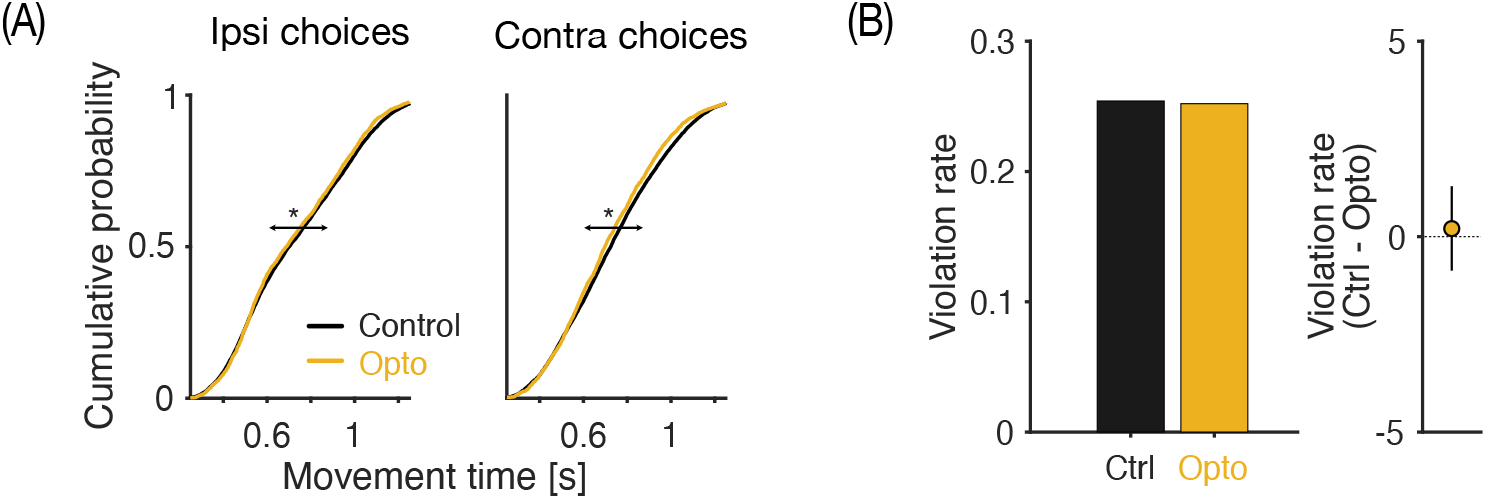
Silencing FOF axon terminals in ADS: effect on movement. **(A)** Silencing reduces movement times. Cumulative distribution of movement times across (n = 6) rats on control (black) and whole-trial inactivation (orange) trials. Movement time is defined as the time taken by the rats to leave the center port and enter one of the two side ports to report their choice. Silencing of FOF axon terminals in ADS significantly reduced the mean movement times for both choices ipsiversive (left, P = 0.05 Mann-Whitney U test) and contraversive (right, P = 0.001 Mann-Whitney U test) to the laser. **(B)** Silencing doesn’t affect rate of trial completion (left) Mean violation rates observed across rats (n=6) on control (black) and whole-trial inactivation (orange) trials. Rats are expected to “fixate” at the center port for 1.5s from trial initiation, failure to do so aborts the trial. Violation rates measure the rate of premature exits from the center port. An increased violation rate might reflect an inability to complete trials due to gross motor/cognitive impairment induced by laser. (right) Mean violation rates between control and opto trial were not different (P = 0.58, non-parametric bootstrap test).

**Supplementary Figure 6.**
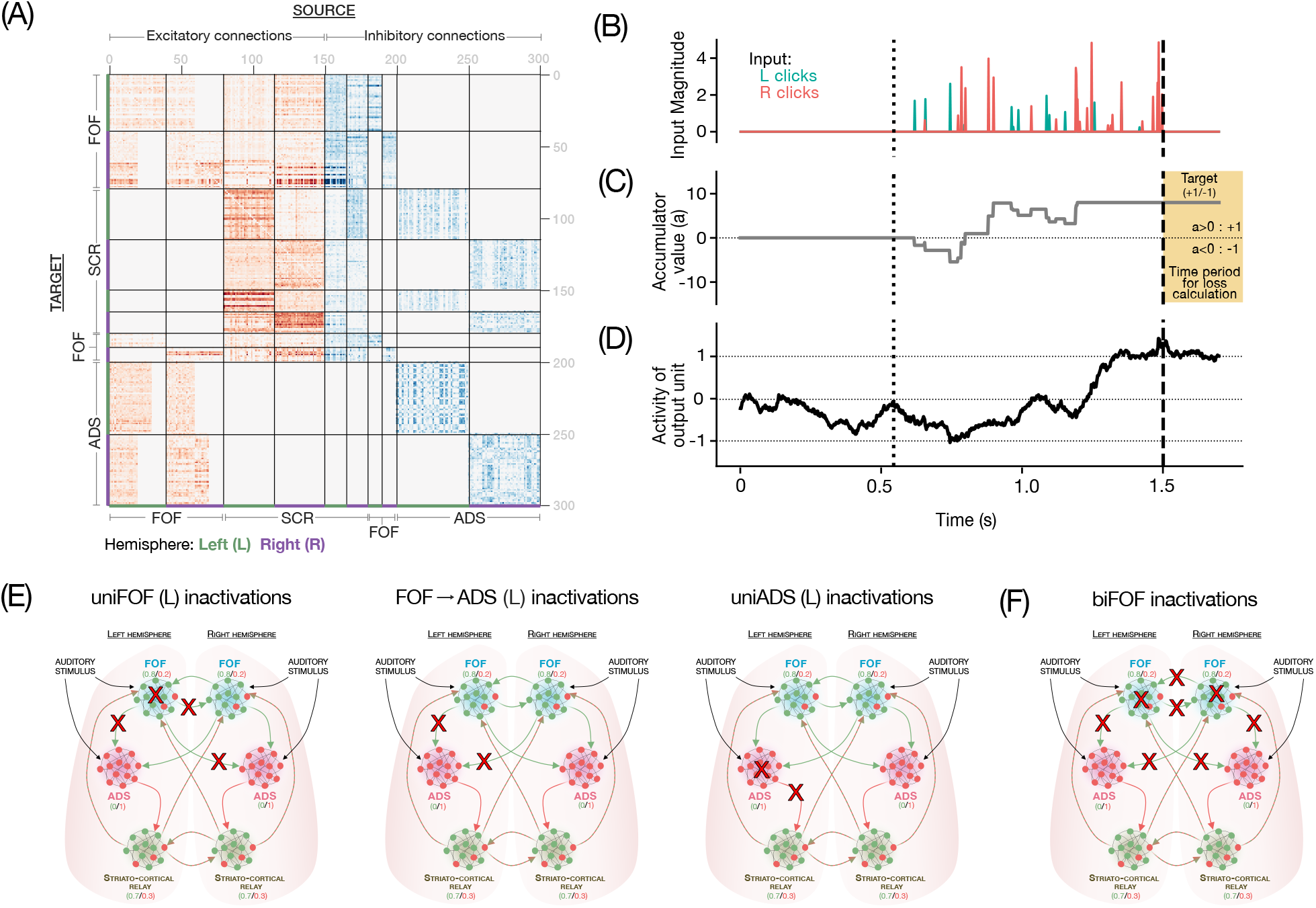
Training details of multi-region recurrent neural network model. **(A)**: Connectivity diagram of recurrent neural network, showing modular structure and following Dale’s law. Excitatory connections are shown in red, and inhibitory in blue, with lines demarcating individual modules in a hemisphere. **(B)**: Example trace of stimulus inputs to the network, showing leftward and rightward clicks with variable timing and magnitudes. **(C)**: Accumulator value computed from stimulus trace above^32^. The final thresholded accumulator value acts as the training target for network outputs on this trial. **(D)**: Example activity of a network output unit that has successfully matched the target output on this trial. **(E)**: Schematics showing all activity and projections that were silenced in unilateral inactivation experiments during training. (Left to Right): unilateral FOF inactivations, FOF → ADS inactivations, unilateral ADS inactivations. Red crosses indicate inactivated elements. **F**: Same as E, but for bilateral FOF inactivations.

**Supplementary Figure 7.**
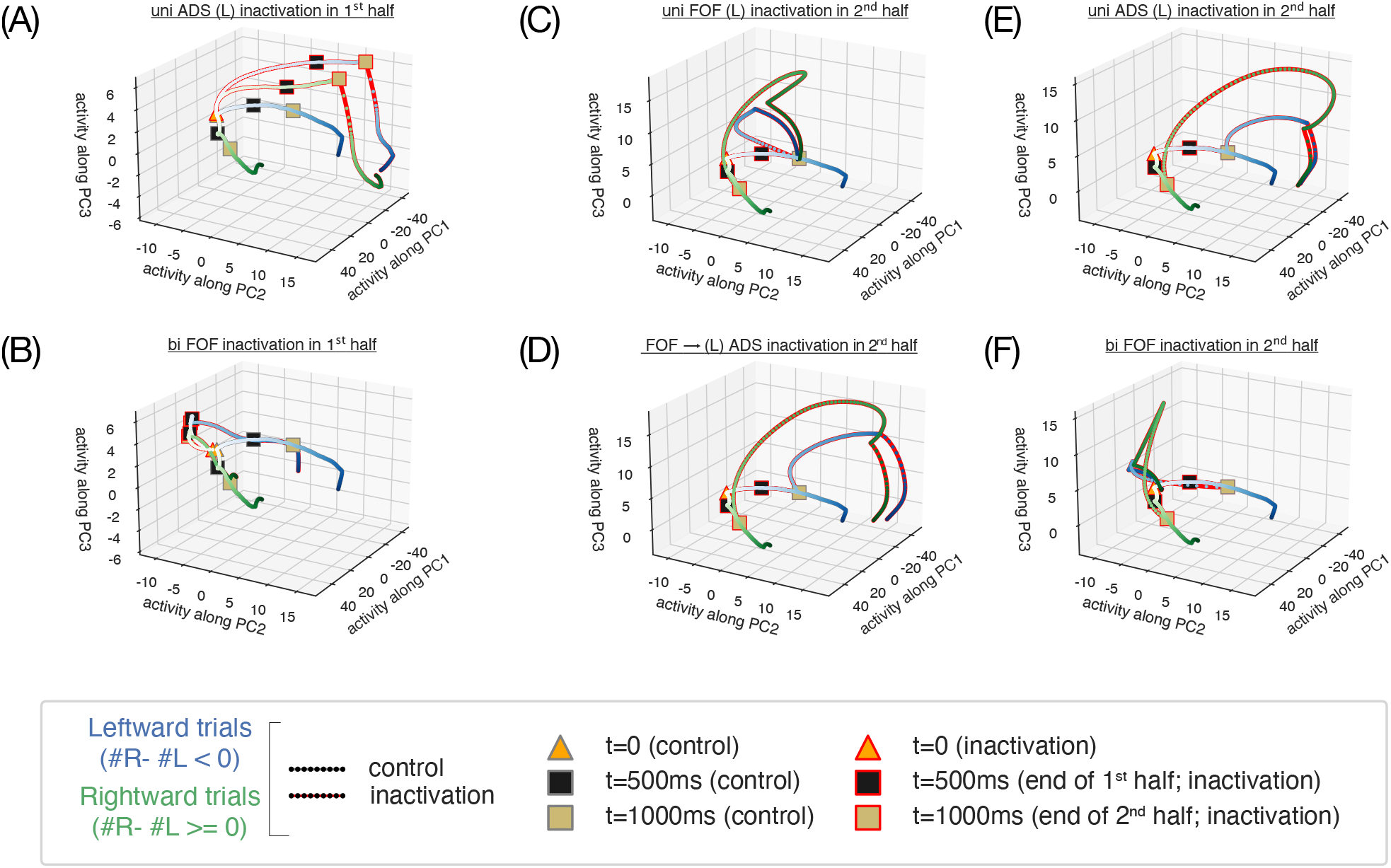
Network dynamics during 1st and 2nd half inactivations. Projections of network activity onto first 3 principal components showing trajectories on rightward (green) and leftward (blue) trials. Red edges represent inactivation trials, and black/yellow boxes represent end of 1st/2nd half respectively. Unilateral inactivation of ADS in the 1st half (A) leads to rightwards trajectories diverging from control and incorrectly driving leftward choices, while bilateral inactivation of FOF in the 1st half (B) recovers towards correct choices subsequently. Unilateral inactivation of all regions in the second half (FOF (C), FOF→ADS projections (D), ADS (E) or bilateral inactivations of FOF (F)) drives incorrect choices and shows no recovery.

**Supplementary Figure 8.**
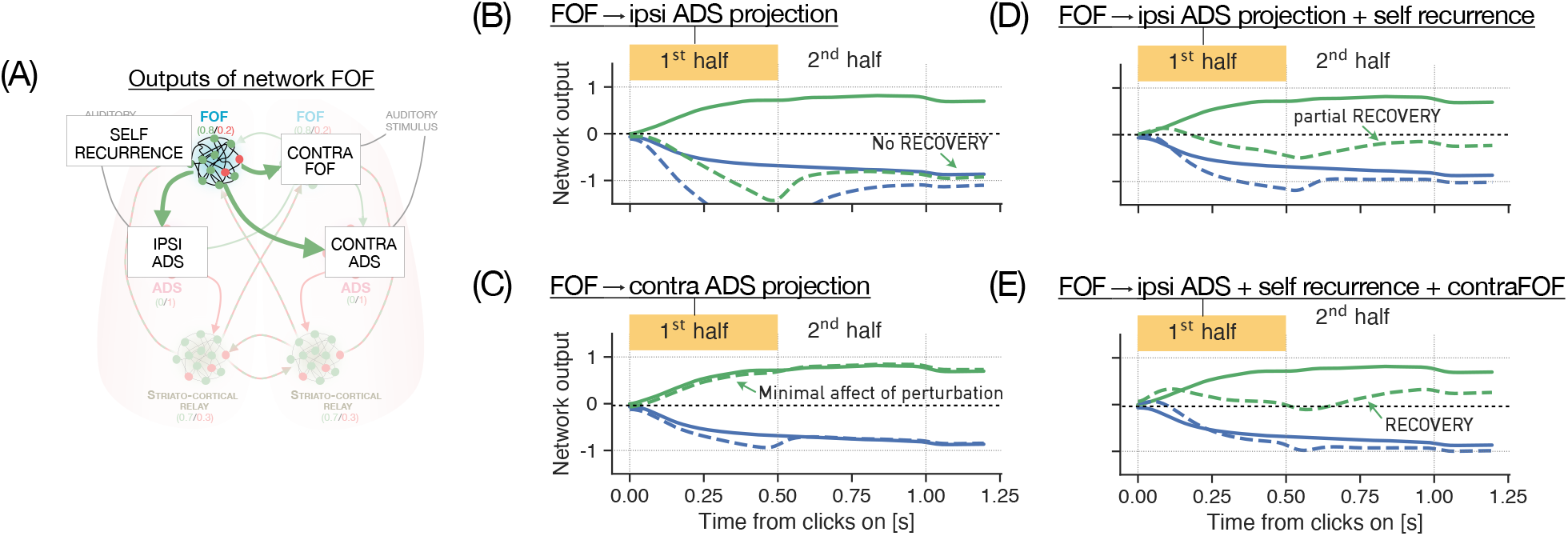
Network outputs in response to in-silico perturbations. **(A)** Schematic of model FOF output projections from a single hemisphere, targeted for in-silico perturbations in the 1st half. Perturbations of left FOF’s projections to ipsilateral ADS (**B**) led to network outputs diverging from control on rightward inactivation trials (dotted green) with no recovery, while perturbations of the projections to contralateral ADS (**C**) had no inactivation effect to begin with - suggesting that the ipsilateral ADS projections were largely responsible for inactivation effects. Additionally perturbing the self recurrence projection (**D**) or the contralateral FOF projection (not shown) invites partial recovery, while perturbing both (**E**) invites nearly complete recovery, with outputs crossing the decision threshold (y=0 line).

